## Supplementary Material for "Beyond SNP Heritability: Polygenicity and Discoverability of Phenotypes Estimated with a Univariate Gaussian Mixture Model"

Holland et al.

#### Relation to Other Work

There is similarity between our work (originally made available on bioRxiv in May 2017 [20]) and the reports of others. Zhu and Stephens [91] implemented a similar causal mixture distribution, but did not take the possibility of inflation into account. They use a reference panel to estimate LD between SNPs, but restrict it to the specific GWAS SNPs, i.e., the ones for which summary statistics are available. Thus, for height, for example, their causal $\beta$ values perforce are restricted to the GWAS SNPs ($\sim$1 million), whereas we use a much larger reference panel ($\sim$11 million) and allow the z-scores in the GWAS of interest to arise from any of the reference panel SNPs. They take care to allow for correlated "noise" in the estimated GWAS effect sizes. Their maximization scheme for the likelihood function involves Markov chain Monte Carlo, and is therefore computationally intensive.

The "M2" model of Zhang et al [14] also uses a causal mixture Gaussian similar to ours. They employ a reference panel of $\sim$1 million SNPs, and allow the causal SNPs to be any of those (thus, in principle, causal SNPs are not necessarily restricted to being GWAS SNPs with z-scores); however, for GWASs with summary statistics for SNPs not in the reference panel, they must restrict the z-scores analyzed to be from SNPs in the reference panel. (In our case, the reference panel is sufficiently large that no such restriction is required.) Their analysis is based on an expression (Eq. (2) in the Supplementary Note of [14], hereinafter "SuppNoteZhang") for the effective $\beta$ at a focal SNP arising through LD from underlying causal $\beta$ values at neighboring SNPs that does not take into account the heterozygosity of those neighboring SNPs – $H_j$ in Eq. (15), $H_1, .., H_w$ in Eq. (20), and $H_w$ in Eq. (26). Their likelihood function (see Eqns. (5) and (7) in SuppNoteZhang, with $H = 1$) ignores the detailed LD and heterozygosity *structure* of the focal SNP (each of the potential causal SNPs in LD with a typed SNP is approximated as having LD with the typed SNP given by the mean for all of the tagged SNPs – the LD score of the typed SNP divided by the number of the SNPs in LD – and their heterozygosities are implicitly treated in an average way, arising through the measured standard error of the $\hat{\beta}$ effect size). We describe how to implement SNP heterozygosity and LD structure in a controlled and accurate way using, equivalently, either Fourier methods or a multinomial expansion (where we take $w = 20$ in our "Model PDF: Multinomial Expansion" subsection, Zhang et al take $w = 1$: compare Eq. (19) here with Eq. (5) in SuppNoteZhang with $H = 1$, and Eq. (21) here with Eq. (7) in SuppNoteZhang, again with $H = 1$; setting $w = 1$ reduces the correct multinomial expansion to an inaccurate binomial expansion). As a result, the M2 model solutions of Zhang et al do not accurately reflect the capability of the underlying mixture distribution for causal effects; their "M3" model, which adds an extra Gaussian, is implemented in the same fashion. The inaccuracy in the M2 model can be seen in the QQ plots (even though they are fairly restricted, with the max value for -$\log_{10}(p)$ set to 10) for, e.g., height, LDL cholesterol, total cholesterol, years of schooling, Crohn's disease, coronary artery disease, and ulcerative colitis, for which we obtain much better fits.

To test the numerical inaccuracy of using the binomial approximation, in simulations, we set $w = 1$ for all cases analyzed in Table 2 and estimated the model M2 parameters from the reulting binomial expansion for the posterior distribution for a SNP's z-score, given the SNP's total LD and heterozygosity. That is, our modified cost function was based on Eq. (21) with $w = 1$. The results, directly comparable with Table 2, are shown below in Table S6. In all cases there is significant underestimation of the M2 heritability, arising from underestimation of the M2 polygenicity or the M2 discoverability, or both. This establishes numerically the mathematical fact mentioned above that the correct multinomial distribution is not equal to the merely approximate binomial distribution.

Table 3 in SuppNoteZhang reports simulations based on "individual level data" (although it is not specified exactly how that was done; in Table 2 they report results from a simulation scheme to generate summary-level association statistics for GWAS without generating individual level data). The closest comparison with our implementation of their M2 model is for the 50k sample size reported in Table 3 SuppNoteZhang where M2 is both the true and fitted model. Although the number of causal SNPs is significantly under-estimated (3.92k vs. 5.35k) and the heritability explained per causal SNP – essentially the discoverability – is significantly over-estimated ($7.6 \times 10^{-5}$ vs. $5.6 \times 10^{-5}$), the heritability turns out to be numerically correct (0.3). In Table S6, for $h^2 = 0.4$ and $n_{causal}$=11k, our estimated number of causal SNPs is only half the true value while the discoverability is a little underestimated ($\hat{\sigma}^2_\beta = 1.2 \times 10^{-4}$ vs. $\sigma^2_\beta = 1.7 \times 10^{-4}$), and $\hat{h}^2 = 0.14$ vs $h^2 = 0.4$. The discrepancies between the implementations of the inaccurate M2 model (in [14] and here) might be due to the factor of ten difference in reference panel size, how exactly individual level data simulations were handled in [14], and/or implementation details. We emphasize, however, that with our full correct implementation, our simulation results are in better accord with the simulated truth. For the example given above, we obtain $\hat{n}_{causal}$=11k, $\hat{\sigma}^2_\beta = 1.6 \times 10^{-4}$ and $\hat{h}^2 = 0.39$, with a correspondingly acurate QQ plot – see Table 2 and Figure S3.

We also applied the M2 model (i.e., setting $w = 1$ and using the binomial expansion as described above) to several of the real phenotypes, and compared with the results of Zhang et al. Below we use a vee ( ˇ ) to distinguish numerical quantities estimated from this procedure. We emphasize that we are using a reference panel with $n_{snp} = 11.02$ million SNPs, whereas Zhang et al use a reference panel with only 1.07 million SNPs. For our larger panel, the mean heterozygosity is $\overline{H} = 0.22$, while for the smaller panel it is $\overline{H} = 0.35$; this difference itself might contribute to differences in estimated heritability, which is a derived quantity that in the model is proportional to the mean heterozygosity in the reference panel.

For height (2010), there were approximately 1 million z-scores, the same as for Zhang et al (essentially the HapMap 3 panel). We obtained $\breve{n}_{causal} = 4.6 \times 10^{-3}$, slightly smaller than the value $4.8 \times 10^{-3}$ from our full correct implementation, Table 1, but in agreement with the value reported by Zhang et al (see Table S5). Inflation (or rather deflation) appears to be minor in the case of height 2010: $\breve{\sigma}_0^2 = 0.94$, the same as our value from the full correct implementation, $\sigma_0^2$ in Table 1. Nonetheless, if we do *not* correct the estimated heritability or discoverability for inflation – i.e., do not divide by $\sigma_0^2$ or $\breve{\sigma}_0^2$ (see Eq. (10)), which Zhang et al do not do ("$a$" in Eq. (3) in [14] corresponds to our $\sigma_0^2$, but values were not reported in [14]) – our M2 estimate is $\breve{h}^2 = 0.15$, which is half the value reported by Zhang et al. These heritability estimates correspond to discoverabilities of $\breve{\sigma}_\beta^2 = 1.53 \times 10^{-4}$ for our implementation of M2 and $1.87 \times 10^{-4}$ for Zhang et al (using $\overline{H} = 0.35$ in Eq. (38)). Note that our M2 estimate for $\breve{\sigma}_\beta^2$ is only slightly smaller that the value $1.56 \times 10^{-4}$ we obtain for the full correct implementation when not rescaling by $\sigma_0^2$ – see Table 1. It is possible that the larger value $1.87 \times 10^{-4}$ for Zhang et al is related to their reference panel being less than a tenth the size of ours.

For schizophrenia, our M2 implementation modeled z-scores from 6.29 million SNPs; the implementation of Zhang et al considerably reduced that to 1.07 million z-scores. Our M2 implementation gave $1.04 \times 10^5$ causal SNPs (versus $3.1 \times 10^4$ for our correct implementation); Zhang et al report $1.9 \times 10^4$ causal SNPs. Our M2 inflation measure is $\breve{\sigma}_0^2 = 1.15$, comparable to our value $\sigma_0^2 = 1.14$ reported in Table 1. Our M2 liability-scale heritability is 0.24 (uncorrected for inflation, which happens to be the same as for our full model also when not correcting for inflation), while Zhang et al report a value of 0.29 – see Table S5. The corresponding discoverabilities are $\breve{\sigma}_\beta^2 = 1.93 \times 10^{-5}$ for our implementation of M2, smaller than $6.28 \times 10^{-5}$ from our full model (again, both values not rescaled for inflation), and $4.36 \times 10^{-5}$ for Zhang et al.

In Table S7 we provide results for all phenotypes analyzed here for which M2 results were available, or implicit, in [14]. We also report height (2014), for which, as with BMI (2015), there were no M2 estimates in [14] though there were M3 estimates. Our M2 estimate of the number of causal SNPs for major depressive disorder, bipolar disorder, schizophrenia, and education all appear to be inflated. It would appear that the inaccurate M2 implementation is increasingly unreliable for higher-polygenicity/lover-discoverability traits. The disjuncture with our M2 simulation results, where we consistently find an underestimation of heritability arising from either an underestimation of polygenicity or discoverability, or of both (Table S6), might be due to effects of model misspecification with respect to real phenotypes exacerbated by the inherent inaccuracy of the implementation, particularly when using a very large reference panel. Use of the smaller reference panel in [14] might be fortuitous in this regard.

Mathematically, we have shown that the binomial implementation of the M2 model used by Zhang et al is inaccurate, and we have demonstrated that numerically with simulations. In contrast, simulations with our correct implementation demonstrate much better parameter estimates – see Table 2. In both scenarios we used exactly the same simulated GWAS z-scores and the full reference panel of 11 million SNPs. When comparing with Zhang et al, there is the additional matter that they are using a reference panel one tenth the size of ours, and, for many real phenotypes (the situation with schizophrenia is representative) they exclude the vast majority of data (z-scores) from the analysis.

Exploring potential MAF-dependence on causal effect sizes, using only GWAS summary data, will be a focus of future work. Our concern here is to present the basic model, correctly implemented and analyzed. Zeng et al [15] use a related causal distribution for true effects $\beta$, but with MAF-dependence. Their approach is based on raw genotype data and requires extensive parallel computing. It is restricted to ∼400k SNPs; with summary statistics, we are working with ∼10 million reference panel SNPs. Below we explore possible model misspecification related to MAF-dependence – see Table S1.

### Data Preparation

Sequentially moving through each chromosome in contiguous blocks of 5,000 SNPs, for each SNP in the block we calculated its Pearson $r^2$ correlation coefficients with all SNPs in the central bock itself and with all SNPs in the pair of flanking blocks of size up to 50,000 each. For each SNP we calculated its total LD (TLD), given by the sum of LD $r^2$'s thresholded such that if $r^2 < r^2_{min}$ we set that $r^2$ to zero. For each SNP we also built a histogram giving the numbers of SNPs in $w_{max}$ equally-spaced $r^2$-windows covering the range $r^2_{min} \leq r^2 \leq 1$. These steps were carried out independently for 1000 Genomes phase 3 and for HAPGEN2 (for the latter, we used 1000 simulated samples).

Employing a similar procedure, we also built binary (logical) LD matrices identifying all pairs of SNPs for which LD $r^2 > 0.8$, a liberal threshold for SNPs being "synonymous".

In applying the model to summary statistics, we calculated histograms of TLD and LD block size (using 100 bins in both cases) and ignoring SNPs whose TLD or block size was so large that their frequency was less than a hundredth of the respective histogram peak; typically this amounted to restricting to SNPs for which TLD $\leq 600$ and LD block size $\leq 1,500$. We also ignored summary statistics of SNPs for which MAF $\leq 0.01$.

For schizophrenia, for example, there were 6,610,991 SNPs with finite z-scores out of the 11,015,833 SNPs from the 1000 Genomes reference panel that underlie the model; the genomic control factor for these SNPs was $\lambda_{GC} = 1.466$. Of these, 314,857 were filtered out due to low MAF or very large LD block size. The genomic control factor for the remaining SNPs was $\lambda_{GC} = 1.468$; for the pruned subsets, with $\simeq 1.49 \times 10^6$ SNPs each, it was $\lambda = 1.30$. (Note that genomic control values for pruned data are always lower than for unpruned data.)

Since effect sizes of individual variants on complex phenotypes are tiny, allelic composition of each variant in case and control groups is very similar, resulting in similar LD structure. There will likely be at most only an extremely slight change in the LD structure for different phenotype-defined subgroups within a population. Imprecision in the phenotype-specificity of the reference panel is currently not a limiting factor in GWAS analysis.

**Summary of the multiple binning process**
(1) We bin typed SNPs (SNPs with z-scores) into an H×L grid. Then, for any grid element, (2) we bin the tagged SNPs into fairly fine LD-$r^2$ bins: the range 0.05-1 is divided up into w=20 or so bins.

In the extreme, we can consider each typed SNP individually, and inquire about how its z-score arises from causal $\beta$ values distributed among the reference SNPs it is in LD with (the SNPs tagged by the typed SNP).

The two levels of binning ((1) and (2) above) can be finessed until we get converged results for our three model parameters – and of course it not necessary to go all the way to the extreme scenario. As mentioned in the main text, we ensured the binning in (1) and (2) was fine enough to give converged results.

From (1) above, we have, in a given grid element, selected typed SNPs in a narrow range of H and L. Conceptually, think of these SNPs as being just a single typed SNP. An important point is that we look at the reference SNPs our typed SNP is in LD with: we group those into very narrow LD bins (w=20 bins). Our model is that the typed SNP's z-score arises from noise and possibly some causal effects among the SNPs it tags through LD: a causal SNP might be in any one of those latter bins, there might be more than one causal SNP in any one of those bins, and many of the bins might have causal SNPs, and all of these distinct possible scenarios need to be accounted for. $k_i$ in Eq. 21 is our random integer variable for the number of causal SNPs in bin i. A second important point is that within bin i, regardless of i, the heterozygosities of the binned reference SNPs will be in a very narrow range – effectively, heterozygosity binning of the SNPs tagged by a given typed SNP comes along free with fine-grained LD $r^2$ binning of those tagged SNPs (again, we're conceptually just dealing here with a single typed SNP whose z-score we are trying to predict). But because all SNPs in bin i have very similar heterozygosities, the heterozygosity of any one of them is necessarily properly accounted for: it is given by $H_i$ in Eq. 21, which carries over entirely to the Fourier version of the z-score pdf – see Eq. 26.

### t-Statistic for Pearson correlation coefficient

The sample Pearson correlation coefficient, $\hat{r}$, and simple linear regression slope, $\hat{\beta}$, for the response and explanatory variables in a data set, are proportional and so have the same t-statistic (and p-value), calculated below.

The true (or population) correlation, $r$, between two random variables $X$ and $Y$ is

$$\begin{aligned} r \equiv \mathrm{corr}(X,Y) &= \frac{\mathrm{cov}(X,Y)}{\sqrt{\mathrm{var}(X)}\sqrt{\mathrm{var}(Y)}} \\ &= \mathrm{corr}(\delta Y + \gamma, \beta X + \alpha) \end{aligned} \tag{40}$$

for any constants $\alpha, \beta, \gamma, \delta$. If one linearly models $Y$ as $Y = \beta X + \alpha$, i.e., $y_i = \beta x_i + \alpha + \epsilon_i$ for the i$^{\mathrm{th}}$ instantiation where $\epsilon_i$ is the residual (random error, assumed to follow a zero-centered normal distribution: $\epsilon_i \sim \mathcal{N}(0, \sigma_\epsilon^2)$), then, by minimizing the mean squared error, the population (true) value $\beta$ is estimated as the sample linear regression coefficient $\hat{\beta}$:

$$\begin{aligned} \hat{\beta} &= \frac{\sum_i (x_i - \bar{x})(y_i - \bar{y})}{\sum_i (x_i - \bar{x})^2} \\ &= \mathrm{cov}(g,y)/\mathrm{var}(g). \end{aligned} \tag{41}$$

For centered sample data, with response variable $y$ and explanatory variable $g$ (both vectors over $n$ samples, with means $\bar{y} = \bar{g} = 0$), one therefore has the *sample* equation

$$y = g\hat{\beta} + \hat{\epsilon} \tag{42}$$

where $\hat{\epsilon}$ is the sample estimated residual. The mean sample response (prediction) for a given value of the explanatory variable is therefore

$$\hat{y} = g\hat{\beta}. \tag{43}$$

From Eq. 40, the sample squared correlation for the quantities in Eq. 42 (and 43) is

$$\begin{aligned} \hat{r}^2 = \mathrm{corr}(g,y)^2 &= \mathrm{corr}(\hat{y},y)^2 \\ &= \frac{\mathrm{cov}(g,y)^2}{\mathrm{var}(g)\mathrm{var}(y)} \\ &= \frac{\mathrm{var}(\hat{y})}{\mathrm{var}(y)} \\ &= \frac{\hat{\beta}^2 \mathrm{var}(g)}{\mathrm{var}(y)}. \end{aligned} \tag{44}$$

The sample residual vector is $\hat{\epsilon} = y - \hat{y}$ with mean 0: $\bar{\hat{\epsilon}} = (1/n)\sum_{i=1}^n \hat{\epsilon}_i = 0$. Note that $\mathrm{var}(y) = \sum_{i=1}^n (y_i - \bar{y})^2/(n-1)$, but $\mathrm{var}(\hat{\epsilon}) = \sum_{i=1}^n (\epsilon_i - \bar{\hat{\epsilon}})^2/(n-2)$: linear regression removes two degrees of freedom, the intercept ($\hat{\alpha} = 0$ in this case) and slope ($\hat{\beta}$). Note also that the above set of vars should strictly be $\widehat{\mathrm{var}}$, indicating that

| $h^2$ | $\hat{h}^2$ | $\pi_1$ | $\hat{\pi}_1$ | $\sigma^2_{\beta_0}$ | $\sigma^2_{\beta_{eff}}$ | $\hat{\sigma}^2_\beta$ | $\hat{\sigma}^2_0$ | $n_{causal}$ | $\hat{n}_{causal}$ |
|---|---|---|---|---|---|---|---|---|---|
| 0.1 | 0.12 | 1E-5 | 1.2E-5 | 2.1E-3 | 4.5E-3 | 4.1E-3 | 1.01 | 110 | 135 |
| 0.1 | 0.09 | 1E-4 | 0.8E-4 | 2.1E-4 | 4.5E-4 | 4.8E-4 | 1.01 | 1101 | 835 |
| 0.1 | 0.08 | 1E-3 | 0.7E-3 | 2.2E-5 | 4.7E-5 | 4.7E-5 | 1.02 | 11015 | 7587 |
| 0.1 | 0.09 | 1E-2 | 0.6E-2 | 2.2E-6 | 4.7E-6 | 6.5E-6 | 1.01 | 110158 | 61888 |
| 0.4 | 0.46 | 1E-5 | 2.1E-5 | 9.5E-3 | 2.2E-2 | 9.2E-3 | 1.02 | 110 | 230 |
| 0.4 | 0.48 | 1E-4 | 1.1E-4 | 8.6E-4 | 1.8E-3 | 1.8E-3 | 1.04 | 1101 | 1216 |
| 0.4 | 0.33 | 1E-3 | 0.9E-3 | 8.8E-5 | 1.9E-4 | 1.5E-4 | 1.06 | 11015 | 9967 |
| 0.4 | 0.33 | 1E-2 | 0.9E-2 | 8.8E-6 | 1.9E-5 | 1.5E-5 | 1.06 | 110158 | 103945 |
| 0.7 | 0.90 | 1E-5 | 2.8E-5 | 1.5E-2 | 3.1E-2 | 1.3E-2 | 1.02 | 110 | 313 |
| 0.7 | 0.80 | 1E-4 | 1.2E-4 | 1.5E-3 | 3.3E-3 | 2.8E-3 | 1.06 | 1101 | 1340 |
| 0.7 | 0.62 | 1E-3 | 0.9E-3 | 1.5E-4 | 3.3E-4 | 2.9E-4 | 1.09 | 11015 | 9832 |
| 0.7 | 0.56 | 1E-2 | 0.9E-2 | 1.5E-5 | 3.3E-5 | 2.5E-5 | 1.10 | 110158 | 103393 |

Table S1: Testing model misspecification: single Gaussian with selection parameter, for twelve single setups (three different heritabilities, each with four different polygenicities – a single instantiation in each case, i.e., not averaged over multiple instantiations). True causal SNPs are a randomly selected fraction $\pi_1$ of all reference SNPs, and their $\beta$ values are randomly drawn from Gaussians with variance $H^S\sigma^2_{\beta_0}$, where $H$ is the causal SNP's heterozygosity, $S = -0.5$, and $\sigma^2_{\beta_0}$ is given in the table. Let $W$ denote the mean of $H^S$ over all causal SNPs. Then $\sigma^2_{\beta_{eff}} \equiv W\sigma^2_{\beta_0}$, which is approximately what the model will try to estimate in $\hat{\sigma}^2_\beta$.

they are estimated from the data, but this hat is dropped for ease of notation. Decompose the total variation in $y$ around the mean:

$$\sum_i (y_i - \bar{y})^2 = \sum_i (y_i - \hat{y}_i)^2 + \sum_i (\hat{y}_i - \bar{y})^2 \tag{45}$$

i.e.,

$$(n-1)\text{var}(y) = (n-1)\text{var}(\hat{y}) + (n-2)\text{var}(\hat{\epsilon}). \tag{46}$$

From Eq. 44, $\text{var}(\hat{y}) = \hat{r}^2\text{var}(y)$, hence the sample mean squared error is

$$\hat{\sigma}^2_\epsilon \equiv \text{var}(\hat{\epsilon}) = \text{var}(y)(1-\hat{r}^2)(n-1)/(n-2). \tag{47}$$

Now, the variance of the sample slope (as an estimate of the population or true slope) arises through the variation in $y_i$, given $g_i$ (the $g_i$ are assumed to be correct and constant explanatory variables). Thus, from Eq. 41,

$$\text{var}(\hat{\beta}) \equiv \text{var}(\hat{\beta}|g) = \text{var}\left(\frac{\sum_i g_i y_i}{\sum_i g_i^2}\middle| g_i\right) \equiv \left[\text{se}(\hat{\beta})\right]^2. \tag{48}$$

where se denotes standard error (dropping the hat as with var). Hence,

$$\text{var}(\hat{\beta}) = \frac{\sum_i g_i^2 \text{var}(y_i|g_i)}{\left(\sum_i g_i^2\right)^2}. \tag{49}$$

But from Eq. 42, $\text{var}(y_i|g_i) \equiv \text{var}(\hat{\epsilon}) = \hat{\sigma}^2_\epsilon$, hence

$$\begin{aligned}\text{var}(\hat{\beta}) &= \hat{\sigma}^2_\epsilon / \sum_i g_i^2 \\ &= \frac{\hat{\sigma}^2_\epsilon}{(n-1)\text{var}(g)} \\ &= \frac{\text{var}(y)(1-\hat{r}^2)}{\text{var}(g)(n-2)} \\ &= \frac{\hat{\beta}^2(1-\hat{r}^2)}{\hat{r}^2(n-2)}.\end{aligned} \tag{50}$$

From Eq. 44, and noting that, as with $\hat{\beta}$ in Eq. 48, $\text{var}(\hat{r}) \equiv \text{var}(\hat{r}|g)$, it immediately follows that $\hat{\beta}/\text{se}(\hat{\beta}) = \hat{r}/\text{se}(\hat{r})$. So to construct a confidence interval for $\hat{r}$, or equivalently for $\hat{\beta}$, form the t-statistic

$$\begin{aligned}t &= \hat{r}/\text{se}(\hat{r}) \\ &= \hat{\beta}/\text{se}(\hat{\beta}) \\ &= \hat{r}\frac{\sqrt{n-2}}{\sqrt{1-\hat{r}^2}}.\end{aligned} \tag{51}$$

### Mean Relative Risk

For logistic linear regression coefficient $\beta$, the odds ratio for disease is $\text{OR} = e^\beta$; for a rare disease, this is approximately equal to the genotypic relative risk: $\text{GRR} \simeq \text{OR}$. Since $\mathbb{E}\left[\beta^2\right] = \sigma^2_\beta$, the mean relative risk $\mathbb{E}\left[\text{GRR}\right] \simeq 1 + \sigma^2_\beta/2$. Thus, for schizophrenia for example, the mean relative risk is $\simeq 1.00003$.

| $h^2$ | $\hat{h}^2$ | $\pi_1$ | $\pi_{1eff}$ | $\hat{\pi}_1$ | $\sigma_b^2$ | $\sigma_c^2$ | $\sigma^2_{\beta eff}$ | $\hat{\sigma}^2_\beta$ | $\hat{\sigma}^2_0$ | $n_{causal}$ | $\hat{n}_{causal}$ |
|---|---|---|---|---|---|---|---|---|---|---|---|
| 0.14 | 0.16 | 1E-5 | 2.9E-6 | 1.3E-5 | 2.2E-3 | 2.2E-2 | 4.3E-3 | 5.2E-3 | 1.01 | 110 | 138 |
| 0.10 | 0.09 | 1E-4 | 2.9E-5 | 5.1E-5 | 2.2E-4 | 2.2E-3 | 4.2E-4 | 7.6E-4 | 1.02 | 1101 | 560 |
| 0.10 | 0.08 | 1E-3 | 2.9E-4 | 3.4E-4 | 2.2E-5 | 2.2E-4 | 4.2E-5 | 1.0E-4 | 1.02 | 11015 | 3779 |
| 0.10 | 0.10 | 1E-2 | 2.9E-3 | 3.3E-3 | 2.2E-6 | 2.2E-5 | 4.2E-6 | 1.2E-5 | 1.01 | 110158 | 35964 |
| 0.51 | 0.58 | 1E-5 | 2.9E-6 | 1.7E-5 | 8.8E-3 | 8.8E-2 | 1.7E-2 | 1.4E-2 | 1.01 | 110 | 189 |
| 0.36 | 0.39 | 1E-4 | 2.9E-5 | 7.2E-5 | 8.8E-4 | 8.8E-3 | 1.7E-3 | 2.3E-3 | 1.04 | 1101 | 794 |
| 0.39 | 0.35 | 1E-3 | 2.9E-4 | 4.8E-4 | 8.8E-5 | 8.8E-4 | 1.7E-4 | 3.1E-4 | 1.05 | 11015 | 5250 |
| 0.40 | 0.36 | 1E-2 | 2.9E-3 | 3.1E-3 | 8.8E-6 | 8.8E-5 | 1.7E-5 | 4.8E-5 | 1.06 | 110158 | 34184 |
| 0.75 | 0.90 | 1E-5 | 2.9E-6 | 2.7E-5 | 1.5E-2 | 1.5E-1 | 2.8E-2 | 1.4E-2 | 1.01 | 110 | 293 |
| 0.62 | 0.67 | 1E-4 | 2.9E-5 | 9.5E-5 | 1.5E-3 | 1.5E-2 | 2.9E-3 | 2.9E-3 | 1.05 | 1101 | 1043 |
| 0.71 | 0.67 | 1E-3 | 2.9E-4 | 5.6E-4 | 1.5E-4 | 1.5E-3 | 2.9E-4 | 5.0E-4 | 1.08 | 11015 | 6115 |
| 0.71 | 0.60 | 1E-2 | 2.9E-3 | 3.8E-3 | 1.5E-5 | 1.5E-4 | 3.0E-5 | 6.6E-5 | 1.09 | 110158 | 42235 |

Table S2: Testing model misspecification: double Gaussian for large and small effects, for twelve single setups (three different heritabilities, each with four different polygenicities – a single instantiation in each case, i.e., not averaged over multiple instantiations). True causal SNPs are a randomly selected fraction $\pi_1$ of all reference SNPs, and their $\beta$ values are randomly drawn from Gaussians with variances $\sigma_b^2$ and $\sigma_c^2 = 10\sigma_b^2$, 90% from the former, 10% from the latter. See Eq 56. $\pi_{1eff}$ and $\sigma^2_{\beta eff}$ are weighted means of the polygenicities and variances for each Gaussian – see text.

### Model Misspecification

**Single Gaussian with selection parameter**

If the true $\beta$ effects are distributed in a way that is different from the assumed one, how meaningful are the parameters and their estimated values? In principle, this is an open-ended issue, but an important assessment can nevertheless be made.

Our model assumed that the $\beta$s are drawn from a single Gaussian with constant variance $\sigma^2_\beta$, for a fraction $\pi_1$ of the reference SNPs:

$$\beta \sim \mathcal{N}(0, \sigma^2_\beta). \tag{52}$$

Evidence has been reported indicating that some causal effects are larger for SNPs with lower heterozygosity. This has lead to consideration of

$$\beta \sim \mathcal{N}(0, H^S \sigma^2_{\beta_0}) \tag{53}$$

where $H$ is the heterozygosity of a causal SNP being assigned the effect $\beta$, $S$ is a new parameter, and $\sigma^2_{\beta_0}$ is a constant. $S < 0$ suggests there is selection pressure. $S = -1$ is implicit in LD Score regression; our model has $S = 0$; intermediate values have been reported by others [15].

Here, in a series of additional simulations using HAPGEN as before with a sample size of 100,000, we set $S = -0.5$, generated new $\beta$ values prescribed by Eq. 53, and calculated new GWAS z-scores, for the twelve heritability $\times$ polygenicity scenarios we used previously – see Table 2, but this time with just a single instantiation in each case. We then fit our model to the simulated z-score. The estimated model parameters $\hat{\pi}_1$, $\hat{\sigma}^2_\beta$, and $\hat{\sigma}^2_0$, along with $\hat{h}^2$ and $\hat{n}_{causal}$, are shown in Table S1.

The proportion of variance explained by variant $i$, whose genotype vector (over $N$ samples) is $g_i$, is $q_i^2 = \text{var}(y; g_i) = \beta_i^2 H_i$, where $y$ is the phenotype vector over the samples. Then, the expected value of this contribution to heritability is

$$\begin{aligned}\langle q_i^2 \rangle &= \langle \beta_i^2 \rangle H_i \\ &= H_i^S \sigma^2_{\beta_0} H_i \\ &= H^{S_i+1} \sigma^2_{\beta_0}.\end{aligned} \tag{54}$$

$h^2$ arises from $\pi_1$ of the $n$ SNPs. So, not knowing which of the SNPs are causal, $h^2$ will approximately be given by

$$h^2 = \sigma^2_{\beta_0} \sum_{i=1}^{n_{causal}} H_i^{S+1} \simeq \pi_1 \sigma^2_{\beta_0} \sum_{i=1}^{n} H_i^{S+1}. \tag{55}$$

We expect that the contribution of $H^S$ to the variances will be absorbed in an average sense into our single variance estimate, $\hat{\sigma}^2_\beta$. Let $W$ denote the mean of $H^S$ over all causal SNPs. Then $\sigma^2_{\beta_{eff}} \equiv W \sigma^2_{\beta_0}$, which is approximately what the model will try to estimate in $\hat{\sigma}^2_\beta$. As can be seen from Table S1, there is reasonable concordance between the model estimates and the true parameters, with greatest mismatch (factor of 2-3) for the very lowest polygenicity ($\pi_1 = 10^{-5}$) scenarios with higher heritabilities (estimating as a mean response from the full set of variants the actual response arising from a small number of SNPs having large effects). In other words, with pronounced selection pressure ($S = -1/2$), our model estimates for polygenicity (number of causal SNPs), discoverability, and heritability give a reasonable indication of the underlying polygenic architecture.

**Double Gaussian: Large and Small Effects**

If the true non-null $\beta$ effects are distributed with respect to two Gaussians, how well does our single-Gaussian based

| $h^2$ | $\hat{h}^2$ | $\pi_1$ | $\hat{\pi}_1$ | $\sigma^2_\beta$ | $\hat{\sigma}^2_\beta$ | $\hat{\sigma}^2_0$ | $n_{causal}$ | $\hat{n}_{causal}$ |
|---|---|---|---|---|---|---|---|---|
| 0.10 | 0.03 | 3.1E-3 | 4.3E-4 | 1.6E-5 | 3.2E-5 | 1.03 | 34540 | 4722 |
| 0.40 | 0.18 | 3.1E-3 | 6.5E-4 | 6.6E-5 | 1.2E-4 | 1.10 | 34718 | 7112 |
| 0.70 | 0.35 | 3.1E-3 | 9.8E-4 | 1.2E-4 | 1.5E-4 | 1.16 | 34370 | 10840 |

Table S3: Testing model misspecification for three different true heritabilities, $h^2 = 0.1, 0.4, 0.7$, for a fixed overall polygenicity, $\pi_1 = 3.1\times10^{-3}$. True causal SNPs are a randomly selected with prior probability $\pi_{1L}$, which decreases from a maximum of 0.005 at $L = 1$ down to 0 at $L \geq 200$. $\pi_1$ is the fraction of all reference SNPs that are causal; their $\beta$ values are randomly drawn from a Gaussian with variances $\sigma^2_\beta$. See Fig S11.

model perform? It may be the case that the majority of causal effects are distributed with respect to a Gaussian with a particular variance, $\sigma_b^2$, while a minority are distributed with respect to a Gaussian with a much larger variance, $\sigma_c^2$. This reflects situations where a minority of causal SNPs have much larger effects than the majority of causal SNPs. Accordingly, here we examine the case where

$$\beta \sim \pi_1\left((1-p_c)\mathcal{N}(0,\sigma_b^2) + p_c\mathcal{N}(0,\sigma_c^2)\right) + (1-\pi_1)\mathcal{N}(0,0). \quad (56)$$

For definiteness, we set $p_c = 0.1$ and $\sigma_c^2 = 10\sigma_b^2$. As with Table S1, we examined twelve scenarios ($\pi_1 = \{10^{-5}, 10^{-4}, 10^{-3}, 10^{-2}\}$, and $h^2 \simeq \{0.1, 0.4, 0.7\}$), using HAPGEN-generated simulated genotypes. In the instantiations generated, the true heritability based on the sample is calculated, which depends on the particular set of $\beta$ values and the heterozygosity of the SNPs they are assigned to. In Table S2 we report the results of the estimated parameters, $\hat{\pi}_1, \hat{\sigma}^2_\beta$, and $\hat{\sigma}^2_0$, when the distribution of causal effects is modeled with Eq 3. We also report the estimated heritability, $\hat{h}^2$ and number of causal SNPs, $\hat{n}_{causal}$. Since the "b"and "c" Gaussians in Eq. 56 are being estimated with a single Gaussian ($\beta \sim \mathcal{N}(0,\sigma^2_\beta)$), we also report effective underlying parameter values: $\pi_{1eff}$, the weighted mean of $\pi_1 \times (1-p_c)$ and $\pi_1 \times p_c$, weighted by $\sigma_b$ and $\sigma_c$ used as proxies for relative importance of the two Gaussians; and $\sigma^2_{\beta eff}$, the overall variance of the true causal effects. It can be seen that $\hat{\sigma}^2_\beta$ reasonably tracks with $\hat{\sigma}^2_{\beta eff}$, while for the larger polygenicities, $\hat{\pi}_1$ reasonably tracks with $\pi_{1eff}$, and the estimated heritabilities, $\hat{h}^2$, are close to the true values, $h^2$. Although this example is not an exhaustive search, the results demonstrate the general applicability and robustness of the model, giving polygenicities, discoverabilities, and heritabilities than are good indicators of the underlying truth.

**Total LD-dependent Prior Probability**

We also examined a highly artificial scenario where the prior probability of a SNP being causal, $\pi_{1L}$, is not a constant but dependent on total LD, $L$. Because the model assumes there is no LD-dependence on whether or not a reference SNP is causal, we do not expect the single Gaussian model to provide a good fit. We assumed that the prior probability of being causal, $\pi_{1L}$, decreases linearly from a maximum of $\pi_{1L} = 0.005$ at $L = 1$, down to $\pi_{1L} = 0.0$ for $L \geq 200$. The overall proportion of causal SNPs is still denoted $\pi_1$. Results are shown in Table S3. There is consistent under-estimation of the heritability. The model QQ plots gave only a very poor fit to the simulated p-values. That is, the obvious poorness of the fits (see Fig S11) gave little credence in the estimated parameters.

### GWAS Replication

An issue that commonly arises in GWAS is whether z-scores in discovery GWAS and those in replication GWAS are consistent. In particular, if some variants reach genome-wide significance on one study but are far from approaching that threshold in a replication study, is there some overlooked problem or are the values in fact statistically consistent? Our model allows for making principled assessments of consistency between discovery and replication GWAS, and we carried out such an assessments in a recent GWAS of bipolar disorder [35].

A discovery z-score from sample size $N_d$ is the sum of two random components

$$z_d = \delta_d + \epsilon_d. \quad (57)$$

A replication z-score from sample size $N_r$ similarly is

$$z_r = \delta_r + \epsilon_r \quad (58)$$

where $\delta$ is the genetic fixed effect (causal for the SNP in question, or LD-mediated from one or more neighboring causal SNPs), and $\epsilon$ is the environmental contribution and noise, modeled as a normal distribution, $\mathcal{N}(0,\sigma_0^2)$, with mean 0 and variance $\sigma^2_{0d}$ or $\sigma^2_{0r}$ (both approximately equal to 1 for BIP data). Effect sizes are related by

$$\delta_r = \sqrt{\frac{N_r}{N_d}}\delta_d. \quad (59)$$

The posterior distribution $\text{pdf}(z_r|z_d)$ is the convolution of $\text{pdf}(\delta_d|z_d)$ with $\mathcal{N}(0,\sigma^2_{0r})$, where

$$\text{pdf}(\delta_d|z_d) = \frac{\text{pdf}(z_d|\delta_d)}{\text{pdf}(z_d)}\text{pdf}(\delta_d) \quad (60)$$

which, using Eq. 59, gives $\text{pdf}(\delta_r|z_d)$;

$$\text{pdf}(z_d|\delta_d) = \phi(z_d;\delta_d,\sigma^2_{0d}), \quad (61)$$

where $\phi(z;\delta,\sigma^2)$ is the Gaussian for $z$ with mean $\delta$ and variance $\sigma^2$; pdf($z_d$) and pdf($\delta_d$) are calculated using the Gaussian mixture model – see Eq. 29 and Eq. 32. It should be noted that these probability densities are SNP-specific in that they depend on the SNP's heterozygosity and LD structure, i.e., the distributions of heterozygosity and LD $r^2$ of its neighbors. The convolution can be written as

$$\begin{aligned}\text{pdf}(z_r|z_d) &= \int_{-\infty}^{\infty} \text{pdf}(\sqrt{\frac{N_d}{N_r}}\delta_r|z_d)\phi(z_r-\delta_r;0,\sigma_{0r}^2)d\delta_r \\ &= \int_{-\infty}^{\infty} \text{pdf}(\delta_d|z_d)\phi(z_r-\sqrt{\frac{N_r}{N_d}}\delta_d;0,\sigma_{0r}^2)d\delta_d.\end{aligned} \tag{62}$$

This can be re-expressed using fast Fourier transforms without the need to perform explicit integration. Now,

$$\text{pdf}(z_r|z_d) = \frac{1}{\text{pdf}(z_d)}\int_{-\infty}^{\infty}\{\text{pdf}(\delta_d)\phi(z_d;\delta_d,\sigma_{0d}^2)\times \phi(z_r-\sqrt{\frac{N_r}{N_d}}\delta_d;0,\sigma_{0r}^2)d\delta_d\}. \tag{63}$$

But

$$\phi(z_d;\delta_d,\sigma_{0d}^2)\phi(z_r-\sqrt{\frac{N_r}{N_d}}\delta_d;0,\sigma_{0r}^2) = \frac{1}{\sigma_{0d}\sigma_{0r}2\pi}\exp\left(\frac{-(z_d-\delta_d)^2}{2\sigma_{0d}^2}\right)\exp\left(-\frac{(z_r-\sqrt{\frac{N_r}{N_d}}\delta_d)^2}{2\sigma_{0r}^2}\right). \tag{64}$$

For the purpose of testing compatibility of discovery and replication z-scores, $\sigma_{0d}\simeq\sigma_{0r}\simeq 1$ can always be achieved in practice by estimating the original values from the data using the model described here, rescaling the discovery and replication z-scores by the respective inflation estimate, and then analyzing the compatibility of the rescaled discovery and replication z-scores. Assuming $\sigma_{0d}\simeq\sigma_{0r}$ and writing it as $\sigma_0$, Eq. 64 is

$$\phi(z_d;\delta_d,\sigma_{0d}^2)\phi(z_r-\sqrt{\frac{N_r}{N_d}}\delta_d;0,\sigma_{0r}^2) = B\phi(\delta_d;A,S^2) \tag{65}$$

where

$$A \equiv \frac{z_d+\sqrt{\frac{N_r}{N_d}}z_r}{1+N_r/N_d}, \tag{66}$$

$$\begin{aligned}B &\equiv \frac{1}{\sigma_0^2 2\pi}\exp\left(\frac{-(z_d^2+z_r^2)}{2\sigma_0^2}\right)\exp\left(-\frac{A^2}{2S^2}\right), \\ S &\equiv \frac{\sigma_0}{\sqrt{1+N_r/N_d}}.\end{aligned} \tag{67}$$

Hence,

$$\begin{aligned}\text{pdf}(z_r|z_d) &= \frac{B}{\text{pdf}(z_d)}\int_{-\infty}^{\infty}\text{pdf}(\delta_d)\phi(\delta_d;A,S^2)d\delta_d \\ &= \frac{B}{\text{pdf}(z_d)}\int_{-\infty}^{\infty}\text{pdf}(\delta_d)\phi(\delta_d-A;0,S^2)d\delta_d \\ &= \frac{B}{\text{pdf}(z_d)}\int_{-\infty}^{\infty}\text{pdf}(\delta)\phi(A-\delta;0,S^2)d\delta.\end{aligned} \tag{68}$$

The integral is just a convolution. Hence, letting $\mathcal{F}$ denote the Fourier transform operator, with $\mathcal{F}^{-1}$ its inverse,

$$\text{pdf}(z_r|z_d) = \frac{B}{\text{pdf}(z_d)}\mathcal{F}^{-1}\Big\{\mathcal{F}[\text{pdf}_\delta(\cdot)]\times\mathcal{F}[\phi(\cdot,0,S^2)]\Big\}(A). \tag{69}$$

Using Eqs. 60 and 62, or equivalently Eq. 69, for any $z_d$, pdf($\delta_r|z_d$) and pdf($z_r|z_d$) can be calculated for finely-spaced vectors with elements $\delta_r$ and $z_r$ respectively ranging from $-a_{lim}$ to $a_{lim}$, with, say, $a_{lim}=12$ (wide enough so that, for any $z_d$, the pdfs start at 0, increase as $\delta_r$ or $z_r$ increases, and then ultimately decrease to 0 again with further increases in the 1st argument). See Figures S1 and S2 which were calculated for discovery and replication bipolar disorder GWAS data [35], where the data (SNPs) are divided up into a $4\times4$ grid of heterozygosity (H) $\times$ total LD (TLD).

To assess whether observed replication z-scores, $z_{ro}$, are statistically consistent with the observed discovery z-scores, $z_{do}$, for a given SNP (explicitly taking into account its H and TLD structure) one can calculate the probability of obtaining a replication z-score, $z_r$, that is "more extreme" that the observed value as follows: if $z_{do}>0$, calculate p($z_r<z_{ro}$) by integrating the pdf thus

$$\text{p}(z_r<z_{ro}|z_{do}) = \int_{-a_{lim}}^{z_{ro}}\text{pdf}(z_r|z_{do})dz_r, \tag{70}$$

and if $z_{do}<0$, calculate p($z_r>z_{ro}$) from

$$\text{p}(z_r>z_{ro}|z_{do}) = \int_{z_{ro}}^{a_{lim}}\text{pdf}(z_r|z_{do})dz_r \tag{71}$$

(small values will indicate outliers). This probability was calculated for 623 rs# SNPs with discovery p-values $<9.97\times10^{-5}$, as described in [35]. For example, in the discovery data set PGC2 ($N_{eff}=49,367$, with 20,352 cases), SNP rs9834970 (H=0.49976, TLD=97.42, shown as the left-most green disc in the bottom-left panel in Fig. S1) had z-score $z_{do}=-7.52$ and p-value $p_{do}=5.54\times10^{-14}$. In the independent replication dataset ($N_{eff}=35,240$, with 9,412 cases) it had $z_{ro}=-1.04$ and $p_{ro}=0.2959$. The expected effect size in the replication dataset given the discovery z-score, $E(\delta_r|z_{do})=-4.03$. The probability of obtaining a replication z-score more extreme than the one observed was p($z_r>z_{ro}|z_{do}$) = 0.012. If the threshold for replicating were set at $p_t=0.05$ (corresponding to $z_t=-1.6449$), then the probability for the replication z-score for SNP rs9834970 to reach this threshold is 0.97.

For a null hypothesis that replication z-scores are consistent with discovery z-scores, there are no rejections ($\alpha = 0.05$) for Bonferroni or Benjamini-Hochberg FDR. I.e., the replication data are, based on our model, consistent with the discovery data.

### Additional notes on HDL

LDSR estimates for SNP heritability for HDL cholesterol is 15.8% [84]. Our estimate is 7%. The LDSR estimate appears to be based on the subset of samples genotyped with genome-wide association study arrays, reported as 94k in [48], the same subset we used. Note that [48] also include another 94k samples genotyped with the Metabochip array (and in cases where Metabochip and GWAS array data were available for the same individuals, they used Metabochip data (to ensure that key variants were directly genotyped rather than imputed).

Our procedure is geared at characterizing the causal variants that can best be described in a distributional sense. To that end, we exclude SNPs that are in very large LD blocks (and SNPs with very low MAF). Thus, it is possible that we miss some large effects. However, such large effects are the ones that are most likely to be found; more important, from our stand point, is making principled estimates of the characteristics of the remaining undiscovered causal SNPs in complex traits. Nevertheless, in forthcoming work, we will analyze all available phenotypes with a more complex model, incorporating multiple causal Gaussian, heterozygosity-dependent variance, and total LD dependence of prior probabilities.

The more direct comparison for our work is with [92], not [48] which has twice the sample size. [92] report 95 lipid-associated loci, 59 of which were new at that time. Based on their Supplementary Table 2, of the 95 loci, 47 were associated with HDL: 17 of these were previously found, an additional 3 were new findings in the previously found loci, and the remaining 27 were findings in new loci. A subsequent "conditional association analysis" identified secondary signals at 26 loci, of which 10 loci resulted in 11 SNPs (two from one locus) being associated with HDL. Their text notes that when the additional SNPs for the 26 loci were combined with the lead SNPs from the 95 loci, 12.1% of total variance in HDL was explained in the Framingham Heart Study, to be compared with our estimate of 3.3% of phenotypic variance explained by genome-wide significant SNPs. If all 95 loci lead SNPs, plus all secondary SNPs from 26 of the loci, the were used to estimate the proportion of total variance explained by genome-wide significant SNPs, that could easily lead to the discrepancy.

Additionally, HDL has very large signals from chromosomes 8, 15, and 16. Thus it might be more appropriate to analyze HDL in a manner similar to AD, separating out chromosome 19. We will address this in future work.

#### Bipolar Disorder: Probability density for replication z-scores given PGC2 discovery z-scores [pdf(zr|zd)]

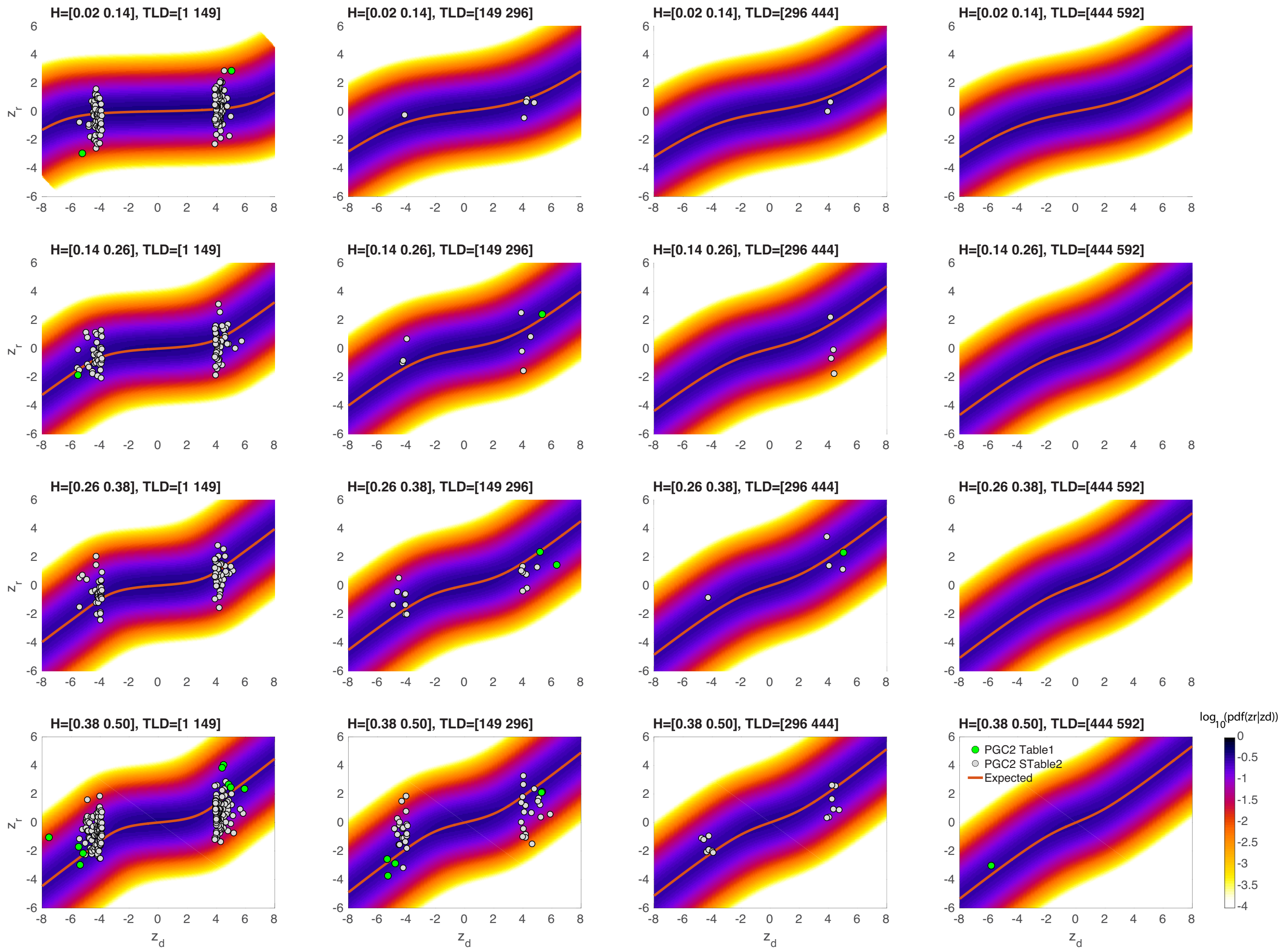


Figure S1: Pdf for replication z-score $z_r$ in a sample with $N_{eff} = 35,240$, given discovery z-score $z_d$ in a sample with $N_{eff} = 49,367$, for bipolar disorder. pdf($z_r|z_d$) is given by Eq. 69. PGC2 Table 1 and PGC2 STable 2 refer to Table 1 and Supplementary STable 2 in [35].

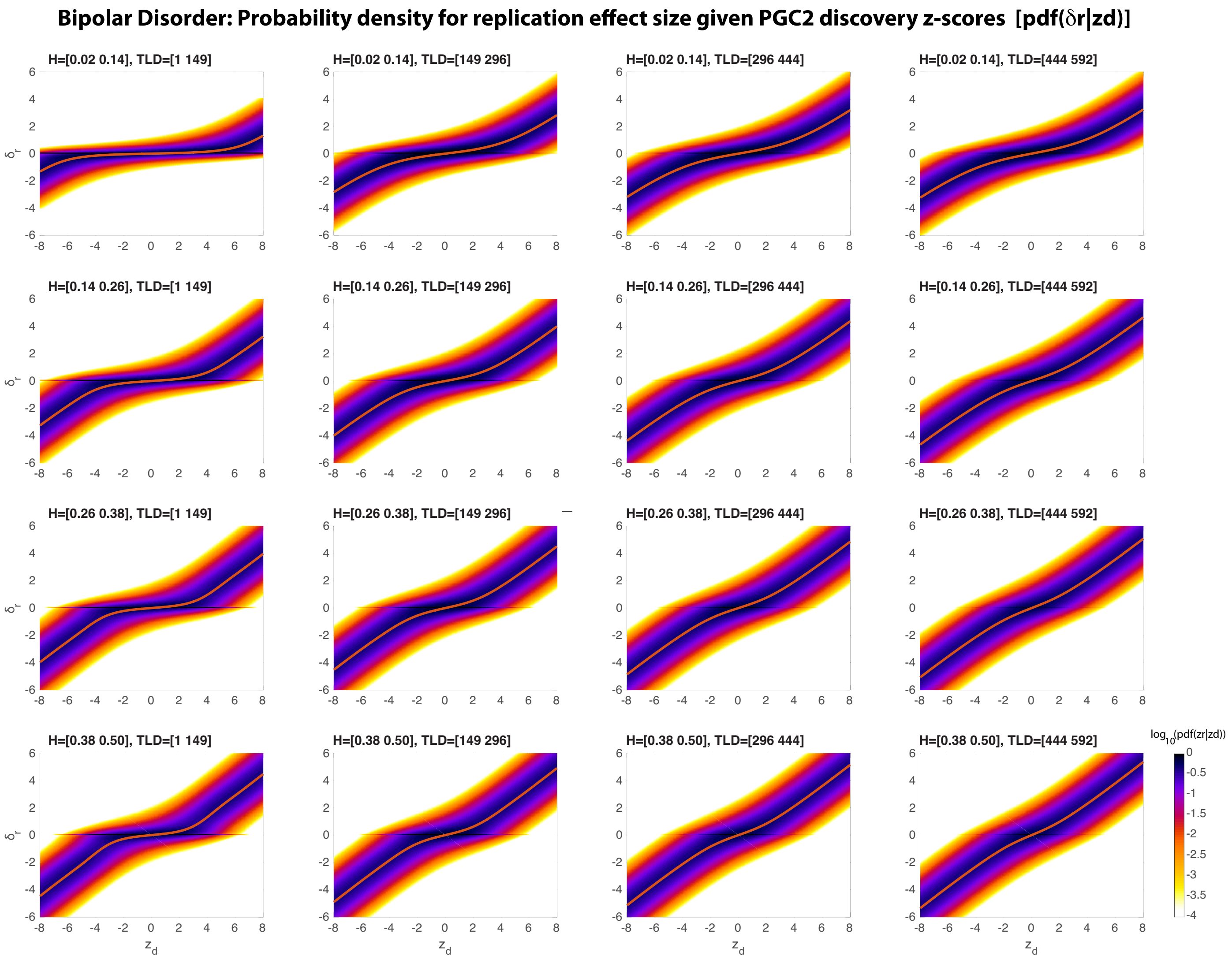


Figure S2: Pdf for replication effect size $\delta_r$ in a sample with $N_{eff} = 35,240$, given discovery z-score $z_d$ in a sample with $N_{eff} = 49,367$, for bipolar disorder. pdf($\delta_r|z_d$) is given by Eqs. 60 and 59.

| Phenotype | $\pi_1$ | $\sigma_b^2$ | $\sigma_0^2$ | $n_{causal}$ | $h^2_{(l)}$ |
|---|---|---|---|---|---|
| MDD | [2.85E-3, 5.17E-3] | [6.21E-6, 8.19E-6] | [1.054, 1.057] | [3.14E4, 5.69E4] | [0.07, 0.07] |
| BIP | [2.46E-3, 2.94E-3] | [4.94E-5, 5.56E-5] | [1.049, 1.051] | [2.71E4, 3.24E4] | [0.16, 0.17] |
| SCZ 2014 | [2.80E-3, 2.87E-3] | [5.47E-5, 5.55E-5] | [1.136, 1.139] | [3.09E4, 3.16E4] | [0.21, 0.21] |
| CAD | [9.94E-5, 1.26E-4] | [1.38E-4, 1.59E-4] | [0.966, 0.973] | [1.09E3, 1.39E3] | [0.03, 0.03] |
| UC | [1.24E-4, 1.28E-4] | [8.71E-4, 8.94E-4] | [1.138, 1.140] | [1.37E3, 1.41E3] | [0.12, 0.12] |
| CD | [9.35E-5, 9.77E-5] | [1.69E-3, 1.71E-3] | [1.166, 1.169] | [1.03E3, 1.08E3] | [0.18, 0.18] |
| AD Chr19 | [7.44E-5, 8.81E-5] | [8.96E-3, 1.72E-2] | [1.066, 1.086] | [1.92E1, 2.28E1] | [0.07, 0.09] |
| AD NoC19 | [1.18E-4, 1.66E-4] | [1.60E-4, 1.97E-4] | [1.044, 1.047] | [1.27E3, 1.79E3] | [0.06, 0.10] |
| ALS Chr9 | [1.24E-5, 1.61E-5] | [0.00, 2.50E-2] | [1.010, 1.021] | [5.13E0, 6.87E0] | [0.00, 0.00] |
| Edu | [3.05E-3, 3.35E-3] | [1.52E-5, 1.62E-5] | [1.003, 1.006] | [3.36E4, 3.69E4] | [0.12, 0.12] |
| IQ 2018 | [2.11E-3, 2.29E-3] | [2.24E-5, 2.39E-5] | [1.279, 1.286] | [2.33E4, 2.53E4] | [0.12, 0.12] |
| BMI | [4.83E-4, 5.00E-4] | [5.47E-5, 5.66E-5] | [0.864, 0.880] | [5.32E3, 5.51E3] | [0.06, 0.07] |
| Height 2010 | [4.13E-4, 4.51E-4] | [1.60E-4, 1.72E-4] | [0.931, 0.940] | [4.55E3, 4.97E3] | [0.17, 0.18] |
| Height 2014 | [5.48E-4, 5.84E-4] | [1.19E-4, 1.27E-4] | [1.656, 1.666] | [6.04E3, 6.43E3] | [0.16, 0.17] |
| Height 2018 | [8.40E-4, 8.71E-4] | [9.29E-5, 9.63E-5] | [2.495, 2.507] | [9.26E3, 9.60E3] | [0.19, 0.20] |
| Putamen | [4.35E-5, 5.53E-5] | [9.27E-4, 1.01E-3] | [1.000, 1.008] | [4.79E2, 6.09E2] | [0.10, 0.12] |
| LDL | [3.31E-5, 3.86E-5] | [5.69E-4, 7.54E-4] | [0.959, 0.965] | [3.63E2, 4.25E2] | [0.05, 0.06] |
| HDL | [2.35E-5, 2.38E-5] | [1.24E-3, 1.26E-3] | [0.969, 0.973] | [2.58E2, 2.62E2] | [0.07, 0.07] |
| TC | [3.96E-5, 4.56E-5] | [8.08E-4, 9.90E-4] | [0.961, 0.966] | [4.36E2, 5.02E2] | [0.09, 0.10] |

Table S4: 95% confidence intervals for the model parameters ($\pi_1$, $\sigma_b^2$ and $\sigma_0^2$), the number of causal SNPs, and the heritability. See Table 1. Confidence intervals for parameters were estimated using the inverse of the observed Fisher information matrix (FIM). The full FIM was estimated for all three parameters used in the model. For the derived quantity $h^2$, which depends on all parameters, the covariances among the parameters, given by the off-diagonal elements of the inverse of the FIM, were incorporated. To calculate the confidence intervals, the mode was run with pruning at $r^2 = 0.1$ to approximate independence of z-scores (for blocks of GWAS SNPs – i.e., SNPs with a z-score – for which their LD $r^2 > 0.1$, a single SNP in the block was randomly chosen to represent the block; blocks were thus approximately independent).

| Phenotype* | $N^{\dagger}_{(eff)}$ | $n_{causal}$ (SE) | $n_{M2}$ (SE) | $n_{M3}$ (SE) | $n_{MS}$ (SE) | $h^2_{(l)}$ (SE) | $h^2_{M2}$ (SE) | $h^2_{M3}$ (SE) | $h^2_{MS}$ (SE) |
|---|---|---|---|---|---|---|---|---|---|
| MDD (2018; 7.1%) | 1.57E5 | 4.4E4 (6.5E3) | 3.0E4 (4.4E3) | — | — | 0.07 (0.001) | 0.09 (0.007) | — | — |
| MDD (UKB; 7.1%) | 2.67E3 | — | — | — | 4.0E4 (1.5E4) | — | — | — | 0.03 (0.005) |
| Bipolar Disorder (0.5%) | 4.94E4 | 3.0E4 (1.3E3) | 1.1E4 (3.5E3) | — | — | 0.16 (0.001) | 0.24 (0.027) | — | — |
| Schizophrenia (1.0%) | 8.07E4 | 3.1E4 (193) | 1.9E4 (1.9E3) | — | — | 0.21 (0.001) | 0.29 (0.013) | — | — |
| CAD (2011; 3.0%) | 6.57E4 | — | 1.9E3 (642) | 2.5E3 (900) | — | — | 0.07 (0.013) | 0.07 (0.012) | — |
| CAD (2015; 3.0%) | 1.63E5 | 1.3E3 (75) | — | — | — | 0.03 (0.001) | — | — | — |
| Ulcerative Colitis (0.1%) | 3.65E4 | 1.4E3 (10) | 856 (321) | 2.7E3 (1.6E3) | — | 0.11 (0.001) | 0.10 (0.015) | 0.13 (0.020) | — |
| Crohn's Disease (0.1%) | 3.62E4 | 1.1E3 (12) | 856 (214) | 6.2E3 (2.1E3) | — | 0.18 (0.001) | 0.17 (0.021) | 0.23 (0.026) | — |
| AD (14.0%) | 4.67E4 | 1.3E3 (140) | 416 (321) | 2.6E3 (1.9E3) | — | 0.15 (0.013) | 0.07 (0.024) | 0.10 (0.021) | — |
| Education (2016) | 2.94E5 | 3.5E4 (837) | 1.9E4 (2.1E3) | — | — | 0.12 (0.001) | 0.13 (0.006) | — | — |
| Education (UKB) | 1.25E5 | — | — | — | 6.4E4 (629) | — | — | — | 0.18 (0.004) |
| Intelligence (2017) | 7.80E4 | — | 1.5E4 (1.9E3) | — | — | — | 0.22 (0.015) | — | — |
| Intelligence (2018) | 2.63E5 | 2.4E4 (512) | — | — | — | 0.13 (0.001) | — | — | — |
| BMI (2010) | 1.24E5 | — | 1.3E4 (1.7E3) | 1.5E4 (1.5E3) | — | — | 0.20 (0.011) | 0.20 (0.010) | — |
| BMI (2015) | 2.34E5 | 7.1E2 (48) | — | 1.8E4 (1.6E3) | — | 0.07 (0.001) | — | 0.13 (0.005) | — |
| BMI (UKB) | 1.26E5 | — | — | — | 4.5E4 (2.4E3) | — | — | — | 0.28 (0.004) |
| Height (2010) | 1.34E5 | 4.8E3 (107) | 4.6E3 (535) | 9.5E3 (1.2E3) | — | 0.17 (0.002) | 0.30 (0.014) | 0.32 (0.015) | — |
| Height (2014) | 2.53E5 | 6.2E3 (100) | — | 1.3E4 (1.3E3) | — | 0.17 (0.001) | — | 0.33 (0.011) | — |
| Height (2018) | 7.08E5 | 9.4E3 (87) | — | — | — | 0.19 (0.001) | — | — | — |
| Height (UKB) | 1.26E5 | — | — | — | 2.3E4 (484) | — | — | — | 0.53 (0.003) |
| Putamen Volume | 1.16E4 | 5.4E2 (33) | — | — | — | 0.11 (0.002) | — | — | — |
| LDL | 8.99E4 | 3.9E2 (16) | 1.4E3 (856) | 9.3E3 (1.9E3) | — | 0.06 (0.002) | 0.08 (0.016) | 0.11 (0.011) | — |
| HDL | 9.43E4 | 2.6E2 (98) | 1.8E3 (963) | 1.0E4 (1.5E3) | — | 0.07 (0.000) | 0.09 (0.015) | 0.11 (0.010) | — |
| Total Cholesterol | 9.46E4 | 4.7E2 (17) | 1.5E3 (535) | 6.4E3 (1.9E3) | — | 0.09 (0.002) | 0.09 (0.013) | 0.12 (0.012) | — |

Table S5: Comparison of our results with those obtained from the two- and three-component Gaussian models in [14], denoted $M2$ and $M3$, respectively, and the single Gaussian model with selection parameter in [15], denoted $MS$. $M2$ and $M3$ use the Hapmap3 reference panel with 1.07 million common SNPs (MAF$\geq$ 0.05); $MS$ uses an Affymetrix panel with 483,634 SNPs (MAF> 0.01) on UK Biobank data; our results are based on a 1000 Genomes Phase3 reference panel with 11 million SNPs (MAF$\geq$ 0.002). SE denotes standard error. $n_{causal}$ is our estimate of the total number of causal SNPs (at MAF>0.2%); $n_{M2}$ and $n_{M3}$ are the total numbers of susceptibility SNPs for $M2$ and $M3$, and $n_{MS}$ the corresponding numbers for $MS$. $h^2_{(l)}$ is our heritability estimate (see Table 1); those obtained from $M2$, $M3$, and $MS$ are labeled $h^2_{M2}$, $h^2_{M3}$, and $h^2_{MS}$, respectively. For quantitative phenotypes, values are on the observed scale; for binary phenotypes, values are on the liability scale, using the same population prevalence as used for Table 1. It is important to note that our heritability estimates are corrected for inflation, by dividing by the inflation parameter $\sigma_0^2$; this is not done for $M2$, $M3$ or $MS$. *Disease prevalences are given as a percentage in parentheses. For binary phenotypes, let $h^2_{obs}$ denote the heritability on the observed 0-1 scale (this is $h^2$ in Figure 1). Let $P$ denote the proportion of cases in the study: $P = N_{cases}/(N_{cases} + N_{controls})$. Then the heritability on the log-odds-ratio scale reported in [14] is $h^2_{log} = h^2_{obs}/(P(1-P))$. The transformation between the observed and liability scale is given by Eq. 39. $^{\dagger}$N is the total sample size for quantitative traits; for qualitative traits, $N_{eff} = 4/(1/N_{cases} + 1/N_{controls})$ – see main text.

| $h^2$ | $\hat{h}^2$ | $\pi_1$ | $\hat{\pi}_1$ | $\sigma^2_\beta$ | $\hat{\sigma}^2_\beta$ | $\hat{\sigma}^2_0$ | $n_{causal}$ | $\hat{n}_{causal}$ |
|---|---|---|---|---|---|---|---|---|
| 0.1 | 0.05 (0.00) | 1E-5 | 8.4E-6 (7E-7) | 4.3E-3 (7E-4) | 2.5E-3 (3E-4) | 1.01 (0.002) | 110 | 92 (7) |
| 0.1 | 0.03 (0.00) | 1E-4 | 4.6E-5 (6E-6) | 4.2E-4 (2E-5) | 3.2E-4 (3E-5) | 1.02 (0.002) | 1101 | 501 (67) |
| 0.1 | 0.04 (0.00) | 1E-3 | 5.2E-4 (8E-5) | 4.2E-5 (5E-7) | 3.3E-5 (3E-6) | 1.01 (0.002) | 11015 | 5714 (903) |
| 0.1 | 0.04 (0.00) | 1E-2 | 2.9E-3 (4E-4) | 4.2E-6 (4E-8) | 5.9E-6 (1E-6) | 1.01 (0.002) | 110158 | 31943 (3950) |
| 0.4 | 0.24 (0.02) | 1E-5 | 1.6E-5 (9E-7) | 1.7E-2 (3E-3) | 6.4E-3 (7E-4) | 1.02 (0.002) | 110 | 172 (10) |
| 0.4 | 0.17 (0.01) | 1E-4 | 6.7E-5 (4E-6) | 1.7E-3 (7E-5) | 1.1E-3 (6E-5) | 1.05 (0.003) | 1101 | 740 (42) |
| 0.4 | 0.14 (0.00) | 1E-3 | 5.0E-4 (3E-5) | 1.7E-4 (2E-6) | 1.2E-4 (6E-6) | 1.06 (0.003) | 11015 | 5510 (359) |
| 0.4 | 0.15 (0.00) | 1E-2 | 3.6E-3 (2E-4) | 1.7E-5 (2E-7) | 1.7E-5 (1E-6) | 1.06 (0.002) | 110158 | 40072 (2572) |
| 0.7 | 0.43 (0.03) | 1E-5 | 2.0E-5 (1E-6) | 3.0E-2 (5E-3) | 9.0E-3 (1E-3) | 1.02 (0.003) | 110 | 220 (14) |
| 0.7 | 0.33 (0.01) | 1E-4 | 8.3E-5 (4E-6) | 2.9E-3 (1E-4) | 1.7E-3 (9E-5) | 1.07 (0.003) | 1101 | 912 (48) |
| 0.7 | 0.25 (0.00) | 1E-3 | 5.1E-4 (2E-5) | 2.9E-4 (4E-6) | 2.0E-4 (6E-6) | 1.10 (0.004) | 11015 | 5661 (241) |
| 0.7 | 0.24 (0.00) | 1E-2 | 3.6E-3 (2E-4) | 2.9E-5 (3E-7) | 2.8E-5 (1E-6) | 1.09 (0.003) | 110158 | 39774 (1775) |

Table S6: Simulation results for our implementation of the M2 model of [14] (i.e., restricting the number of LD $r^2$ windows for the range $0 \leq r^2 \leq 1$ to 1 – see the subsection *Relation to Other Work* on page S2). Shown here is a comparison of mean (std) true and estimated ($\hat{}$) model parameters and derived quantities. Results for each line, for specified heritability $h^2$ and fraction $\pi_1$ of causal SNPs, are from 10 independent instantiations with random selection of the $n_{causal}$ causal SNPs that are assigned a $\beta$-value from the standard normal distribution. Compare with Table 2.

| Phenotype* | $n_{causal}$ | $\check{n}_{causal}$ | $n_{M2}$ | $\sigma^2_\beta$ | $\check{\sigma}^2_\beta$ | $\sigma^2_{M2}$ | $h^2_{(l)}$ | $\check{h}^2_{(l)}$ | $h^2_{M2}$ | $\sigma^2_0$ | $\check{\sigma}^2_0$ |
|---|---|---|---|---|---|---|---|---|---|---|---|
| MDD (2018; 7.1%) | 4.4E4 | 9.8E4 | 3.0E4 | 7.63E-6 | 3.43E-6 | 8.57E-6 | 0.07 | 0.08 | 0.09 | 1.06 | 1.06 |
| Bipolar Disorder (0.5%) | 3.0E4 | 1.5E5 | 1.1E4 | 5.51E-5 | 1.27E-5 | 6.23E-5 | 0.17 | 0.20 | 0.24 | 1.05 | 1.05 |
| Schizophrenia (1.0%) | 3.1E4 | 1.0E5 | 1.9E4 | 6.28E-5 | 1.93E-5 | 4.36E-5 | 0.24 | 0.24 | 0.29 | 1.14 | 1.15 |
| Ulcerative Colitis (0.1%) | 1.4E3 | 1.1E4 | 856 | 1.01E-4 | 1.20E-3 | 4.34E-4 | 0.13 | 0.13 | 0.10 | 1.14 | 1.15 |
| Crohn's Disease (0.1%) | 1.1E3 | 928 | 856 | 1.99E-3 | 2.21E-3 | 5.67E-4 | 0.21 | 0.20 | 0.17 | 1.17 | 1.18 |
| Education (2016) | 3.5E4 | 1.3E5 | 1.9E4 | 1.57E-5 | 4.73E-6 | 1.95E-5 | 0.12 | 0.13 | 0.13 | 1.00 | 1.02 |
| Height (2010) | 4.8E3 | 4.6E3 | 4.6E3 | 1.56E-4 | 1.53e-4 | 1.87e-4 | 0.16 | 0.15 | 0.30 | 0.94 | 0.94 |
| Height (2014) | 6.2E3 | 6.1E3 | — | 2.04E-4 | 1.98E-4 | — | 0.28 | 0.26 | — | 1.66 | 1.66 |
| LDL | 390 | 371 | 1.4E3 | 6.35E-4 | 6.05E-4 | 1.63E-4 | 0.06 | 0.05 | 0.08 | 0.96 | 0.96 |
| HDL | 260 | 255 | 1.8E3 | 1.21E-3 | 1.12E-3 | 1.43E-4 | 0.07 | 0.06 | 0.09 | 0.97 | 0.97 |
| Total Cholesterol | 469 | 290 | 1.5E3 | 8.63E-4 | 6.08E-4 | 1.71E-4 | 0.09 | 0.04 | 0.09 | 0.96 | 0.95 |

Table S7: Comparison of model estimates: (1) our results reported in Table 1 ($n_{causal}$, $\sigma^2_\beta$, $h^2_{(l)}$, and $\sigma^2_0$) without rescaling discoverability and heritability by the inflation parameter; (2) results of our implementation (quantities with a $\check{}$) of the M2 model in [14], without rescaling with respect to $\check{\sigma}^2_0$ – see the first section of this Supplement, *Relation to Other Work*; and (3) M2 results reported or implicit in [14] (quantities with the underscore $_{M2}$). $\sigma^2_{M2} = (h^2_{M2}/n_{M2})/\bar{H}$, where $\bar{H} = 0.35$. *Disease prevalences are given as a percentage in parentheses. See Table S5.

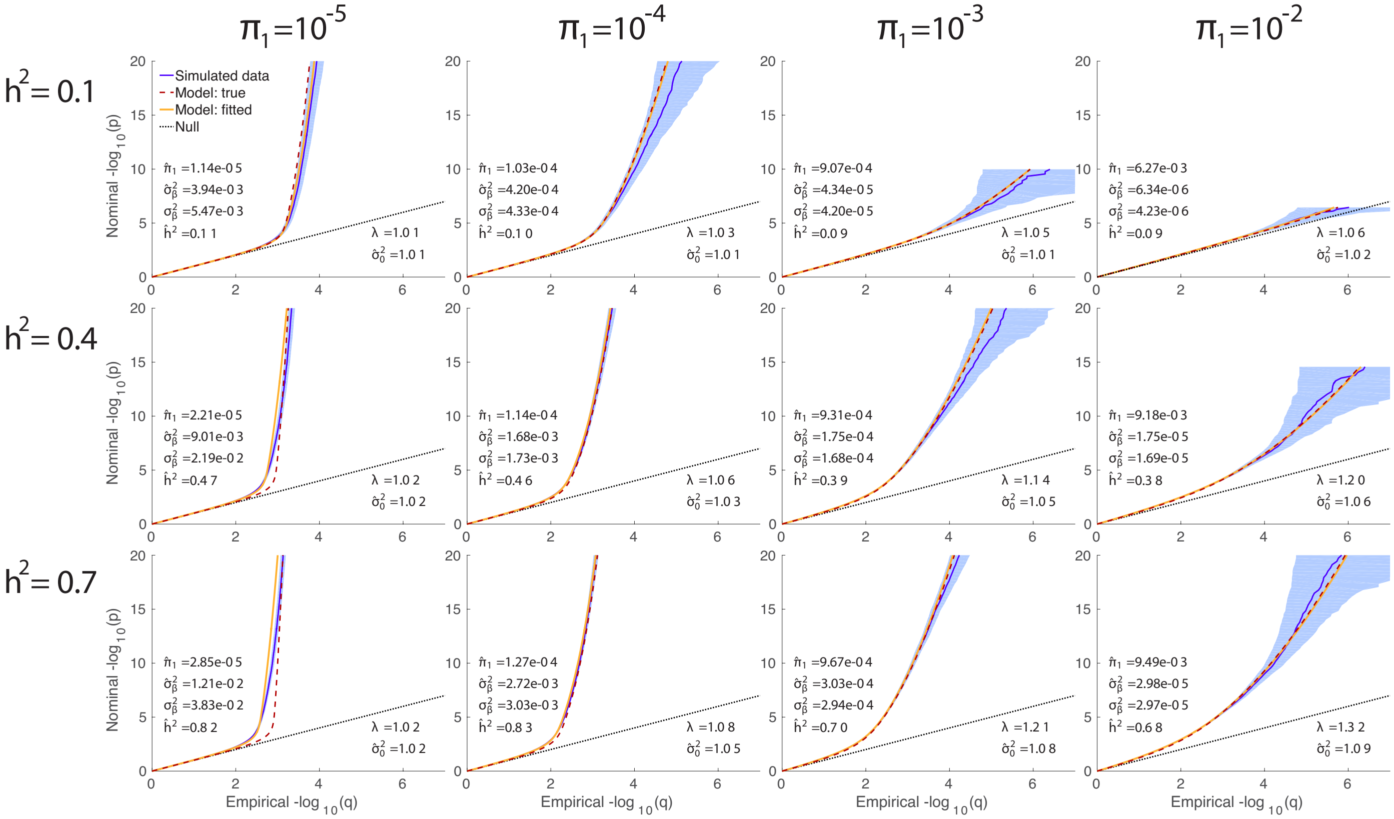


Figure S3: Quantile-quantile plots for simulations. True polygenicity is specified for each column, and true heritability is specified for each row. QQ-plots for simulated data in dark blue, with 95% confidence interval in light blue; model prediction in yellow. The dashed red curve is the QQ plot corresponding to the true parameters. $\lambda$ is the overall nominal genomic control factor for the pruned simulated data. The three estimated model parameters are: polygenicity, $\hat{\pi}_1$; discoverability, $\hat{\sigma}^2_\beta$; and SNP association $\chi^2$-statistic inflation factor, $\hat{\sigma}^2_0$. $\hat{h}^2$ is the estimated narrow-sense chip heritability. The dotted black line is the expected plot under null. Reading the plots: on the vertical axis, choose a p-value threshold (more extreme values are further from the origin), then the horizontal axis gives the proportion of SNPs exceeding that threshold (higher proportions are closer to the origin).

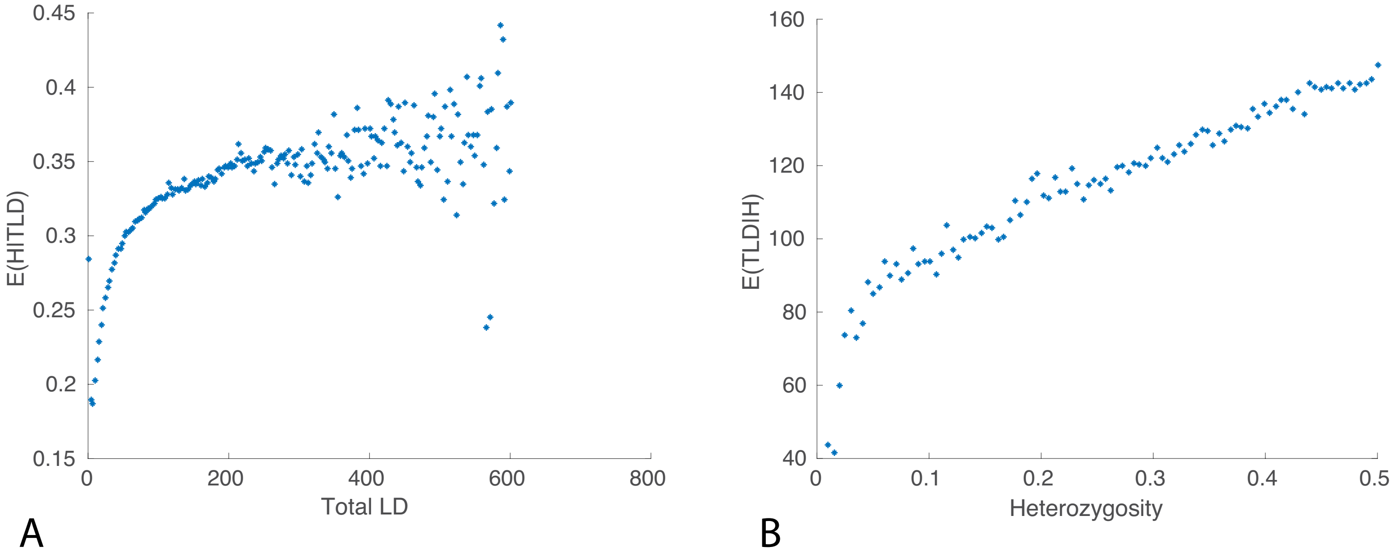


Figure S4: (A) Mean value of heterozygosity for given total LD (SNPs were binned based on TLD and the mean TLD for each bin plotted on the x-axis; the corresponding mean heterozygosity for SNPs in each bin was then plotted on the y-axis). (B) Mean value of total LD for given heterozygosity. Plots made for SNPs in the PGC2 schizophrenia GWAS; TLD and H calculated from 1000 Genomes phase 3 reference panel.

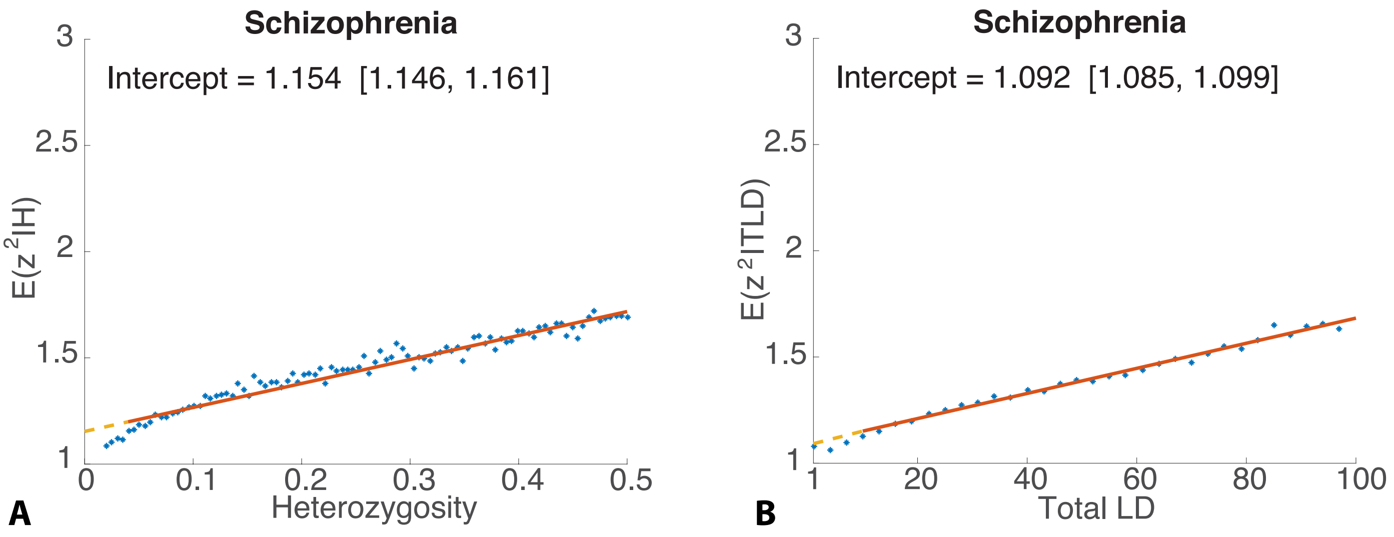


Figure S5: Regressing $z^2$ on (A) heterozygosity, and (B) total LD for schizophrenia [36].

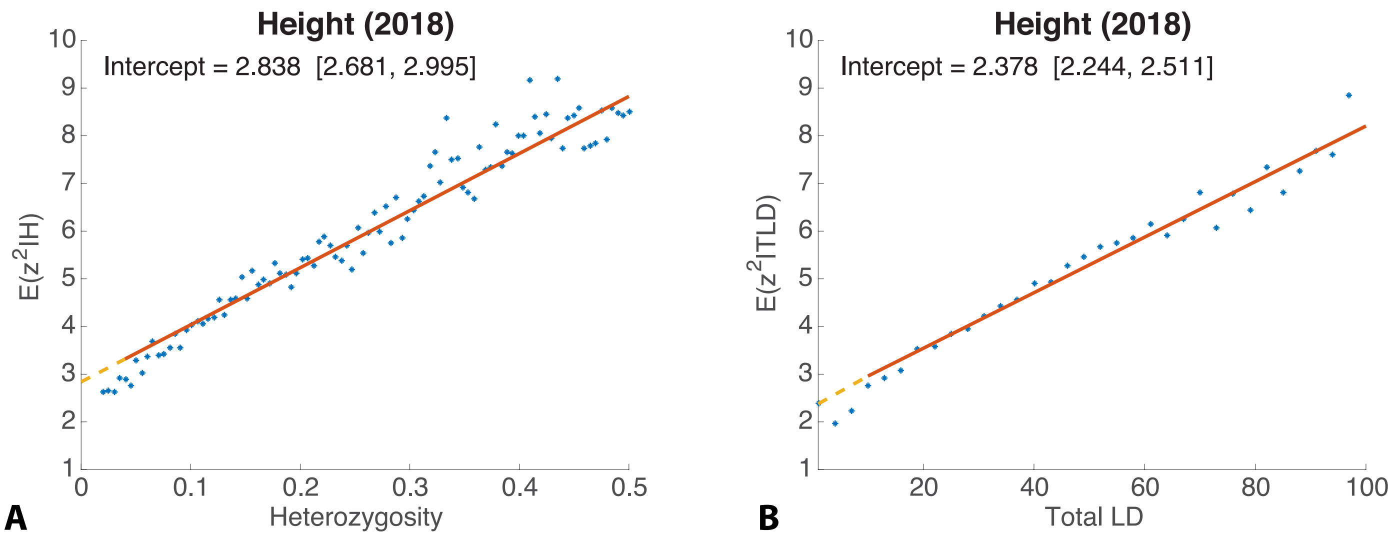


Figure S6: Regressing $z^2$ on (A) heterozygosity, and (B) total LD for height [49].

### Alzheimer's Disease

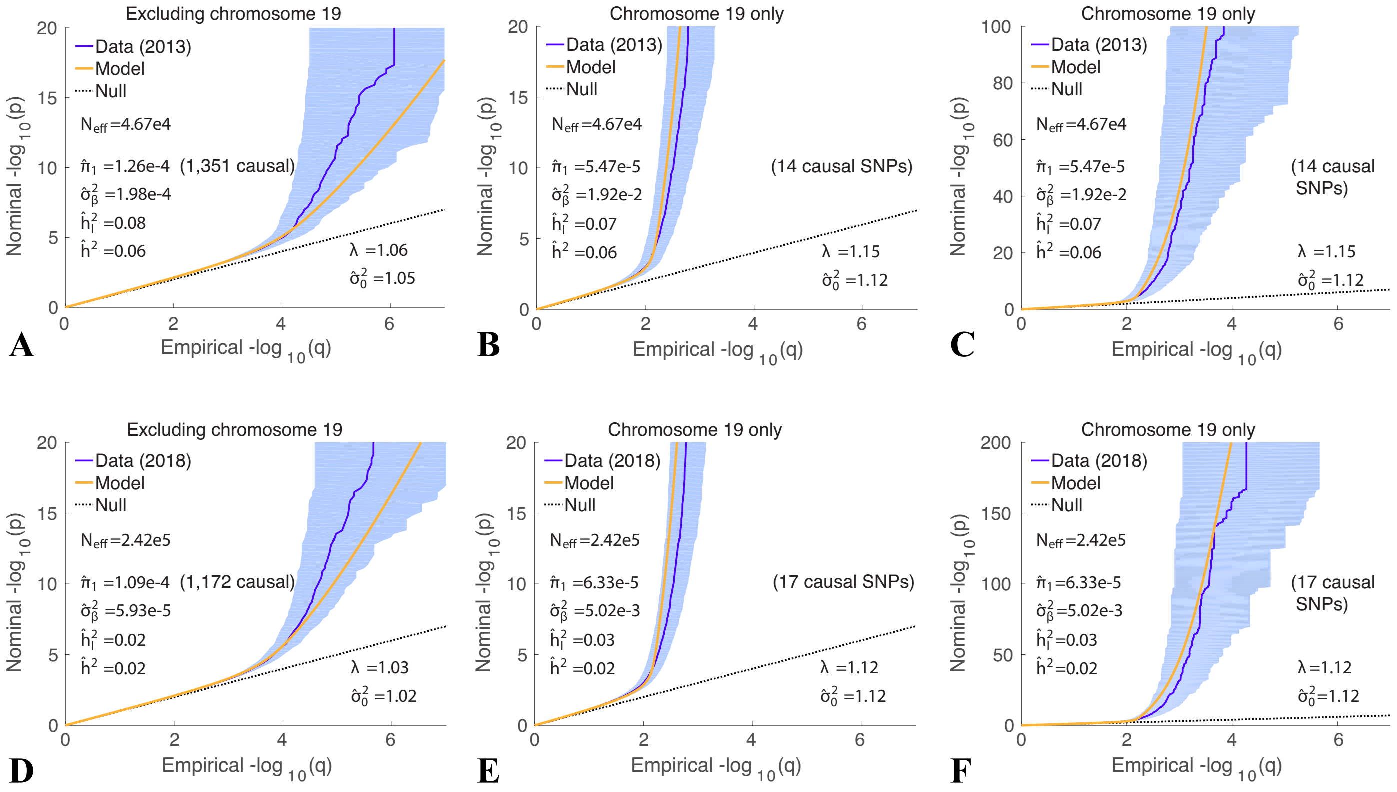


Figure S7: QQ plots for AD, comparing the 2013 data [39] and model results (top row) with those of 2018 [40] (bottom row). For both data sets, we exclude (in (A) and (D)) chromosome 19 from the analysis, and in (B) and (E) we analyze chromosome 19 exclusively. (C) is (B) with the y-axis extended 5×; (F) is (E) with the y-axis extended 10×. The heritabilities reported are for the relevant sections of DNA. From (A) and (B), the total (full autosomal reference panel) narrow-sense liability-scale SNP heritability for AD is $h_l^2 = 0.15$. See Figure 1 for a further description.

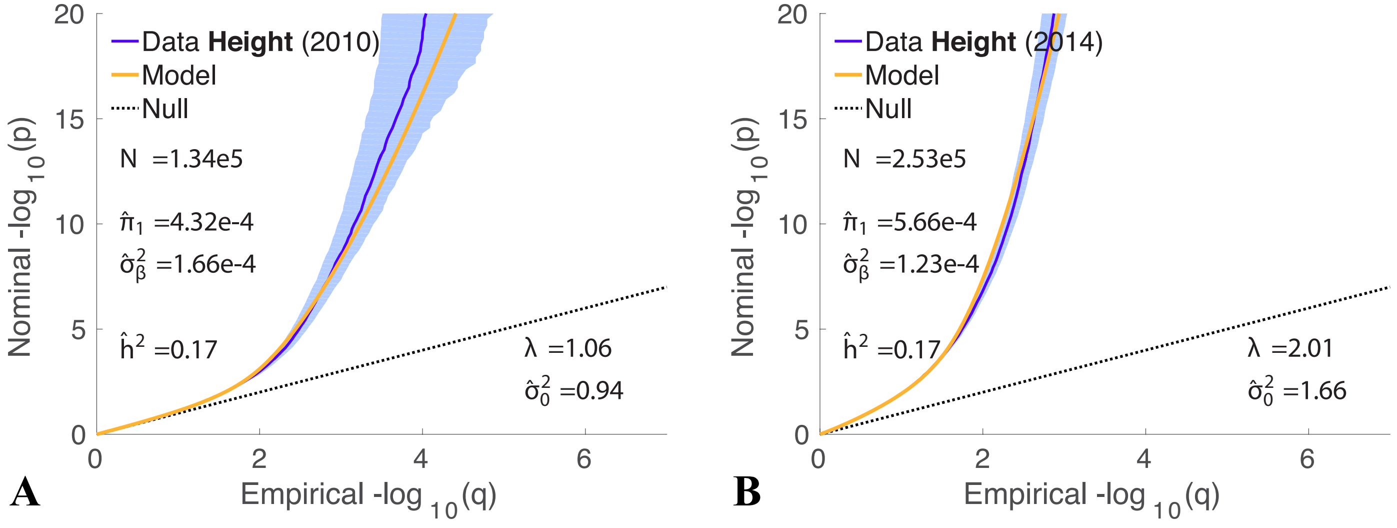


Figure S8: QQ data and model plots for height, comparing the (A) 2010 [65] and (B) 2014 [46] GWAS. See also Figure 2 D, Table 1, and Figure S10.

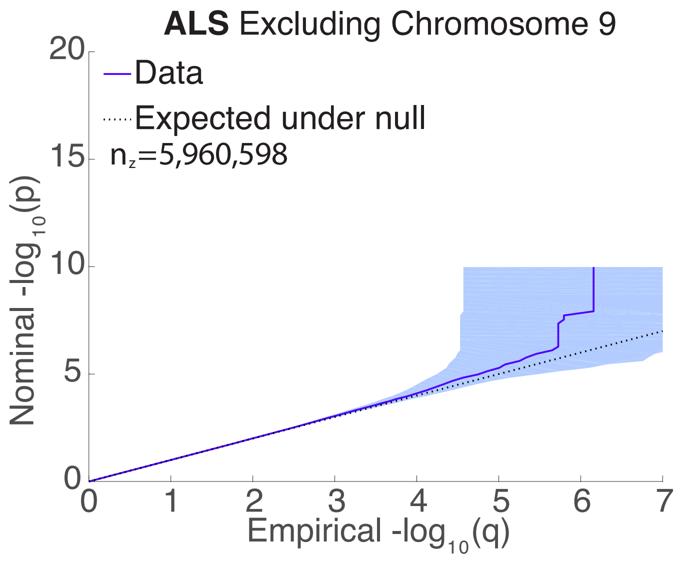


Figure S9: QQ plot for amyotrophic lateral sclerosis (ALS) excluding chromosome 9 (z-scores randomly pruned at correlation $r^2 = 0.8$). The data slightly deviate from the null line only after the proportion (q) is less than one in a million: only seven z-scores outside chromosome 9 reach genome-wide significance. This is an extremely weak signal where the model is not expected to be informative. Compare with Figure 1 (H).

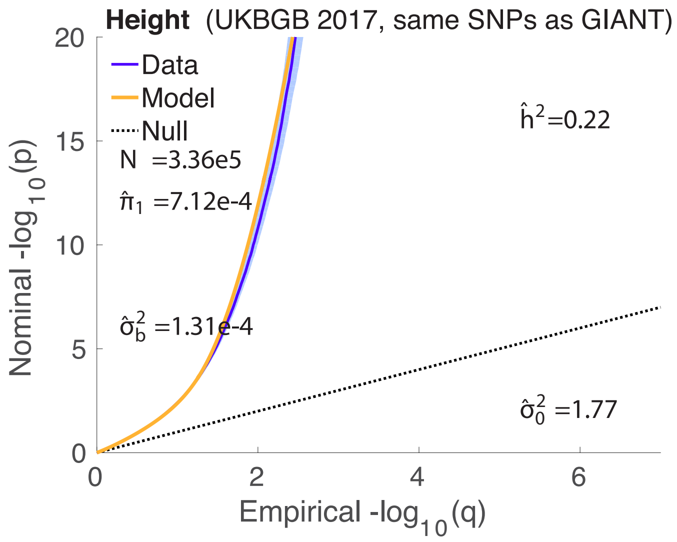


Figure S10: QQ plot for UKB GB height (2017), with the same set of SNPs as in the 2010 and 2014 GIANT datasets. Compare with Figure S8.

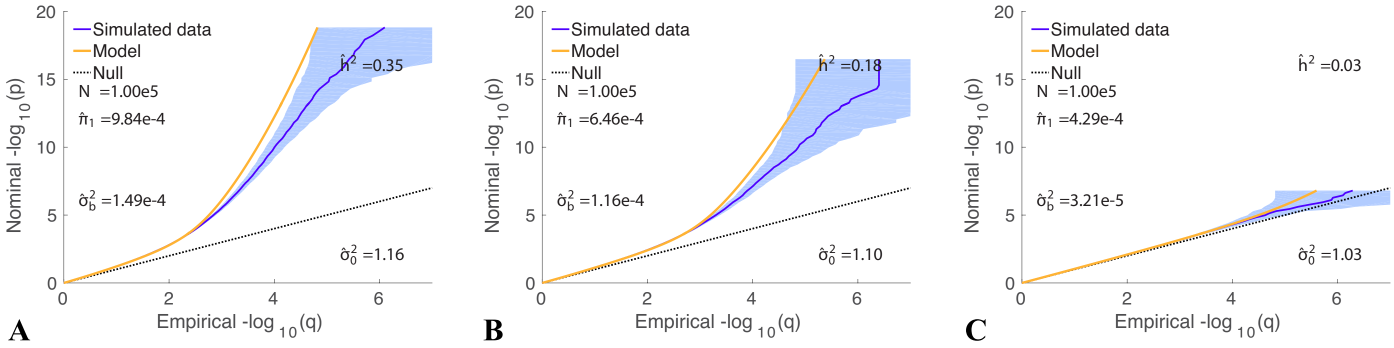


Figure S11: Model misspecification with a fixed overall polygenicity, $\pi_1 = 3.1 \times 10^{-3}$, for three different true heritabilities: (A) $h^2 = 0.7$; (B) $h^2 = 0.4$; and (C) $h^2 = 0.1$. True causal SNPs are a randomly selected with prior probability $\pi_{1L}$, which decreases from a maximum of 0.005 at $L = 1$ down to 0 at $L \geq 200$. $\pi_1$ is the fraction of all reference SNPs that are causal; their $\beta$ values are randomly drawn from a Gaussian with variances $\sigma_\beta^2$. See Table S3.

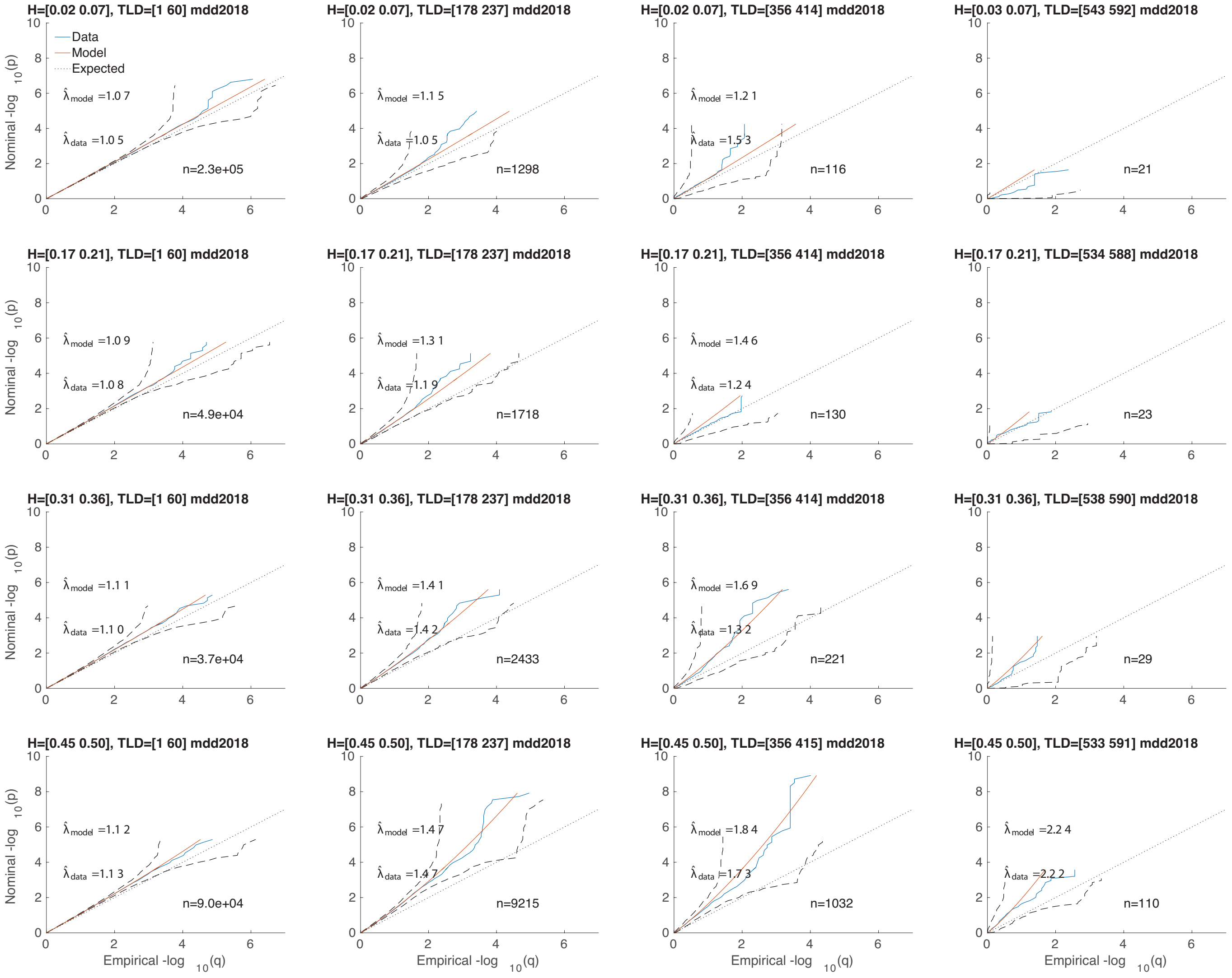


Figure S12: A 4×4 heterozygosity×total-LD grid of QQ plots for major depressive disorder (H increasing top to bottom, TLD increasing left to right, taking every second subplot from a full 10×10 grid). $n$ is the number of SNPs in each grid element. See Figure 1 (A).

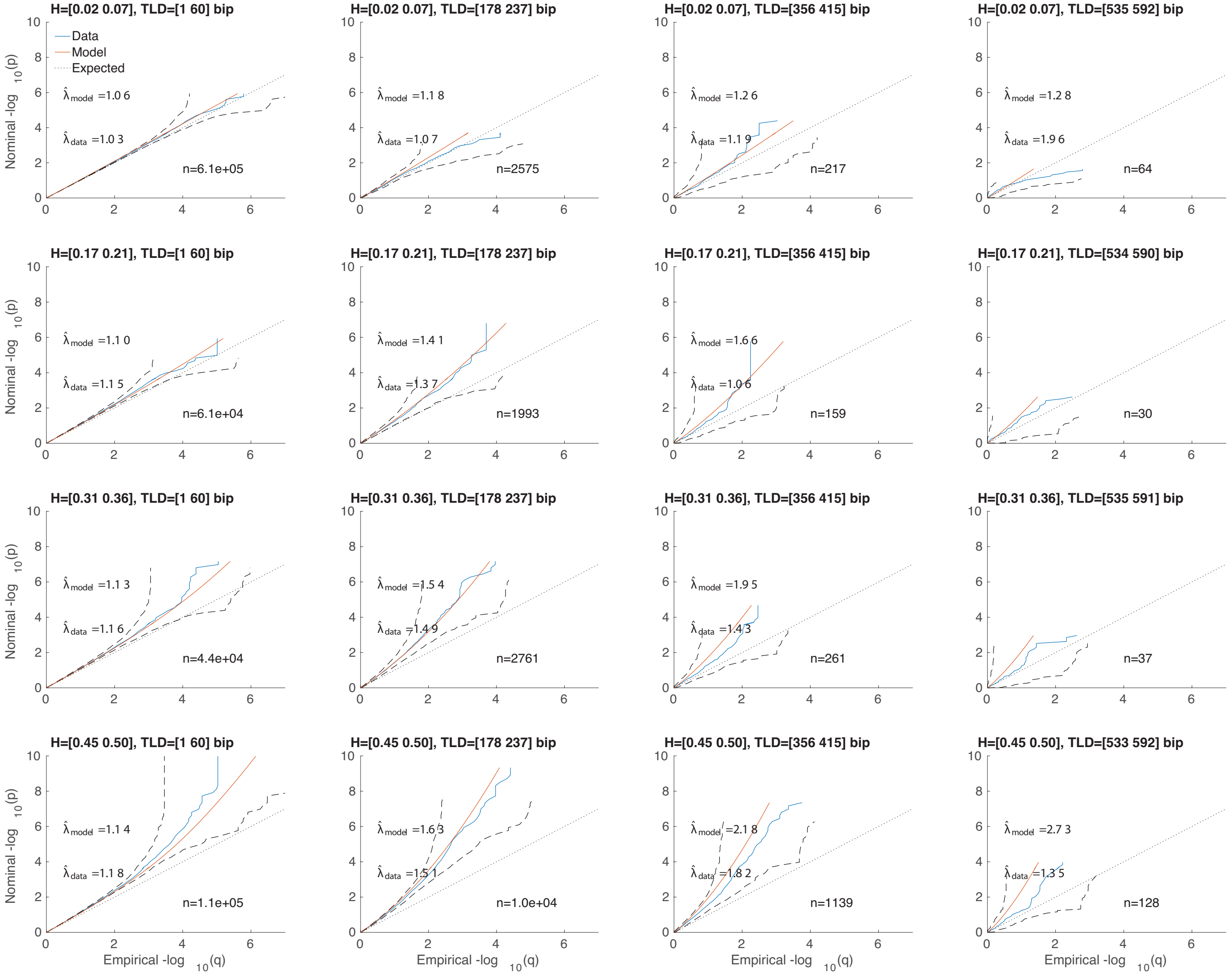


Figure S13: A 4×4 heterozygosity×total-LD grid of QQ plots for bipolar disorder. See Figures 1 (B) and S12.

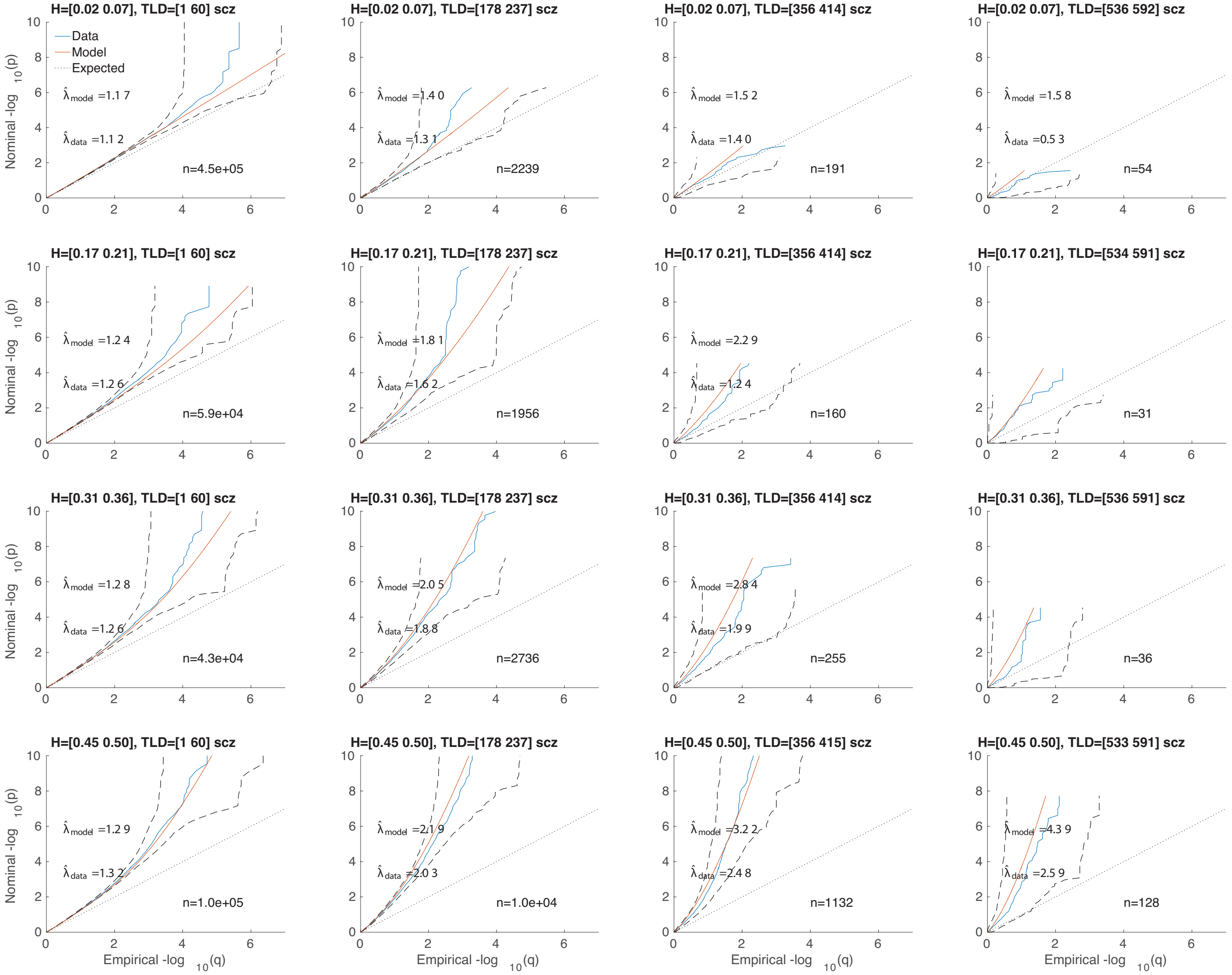


Figure S14: A 4×4 heterozygosity×total-LD grid of QQ plots for schizophrenia. See Figures 1 (C) and S12.

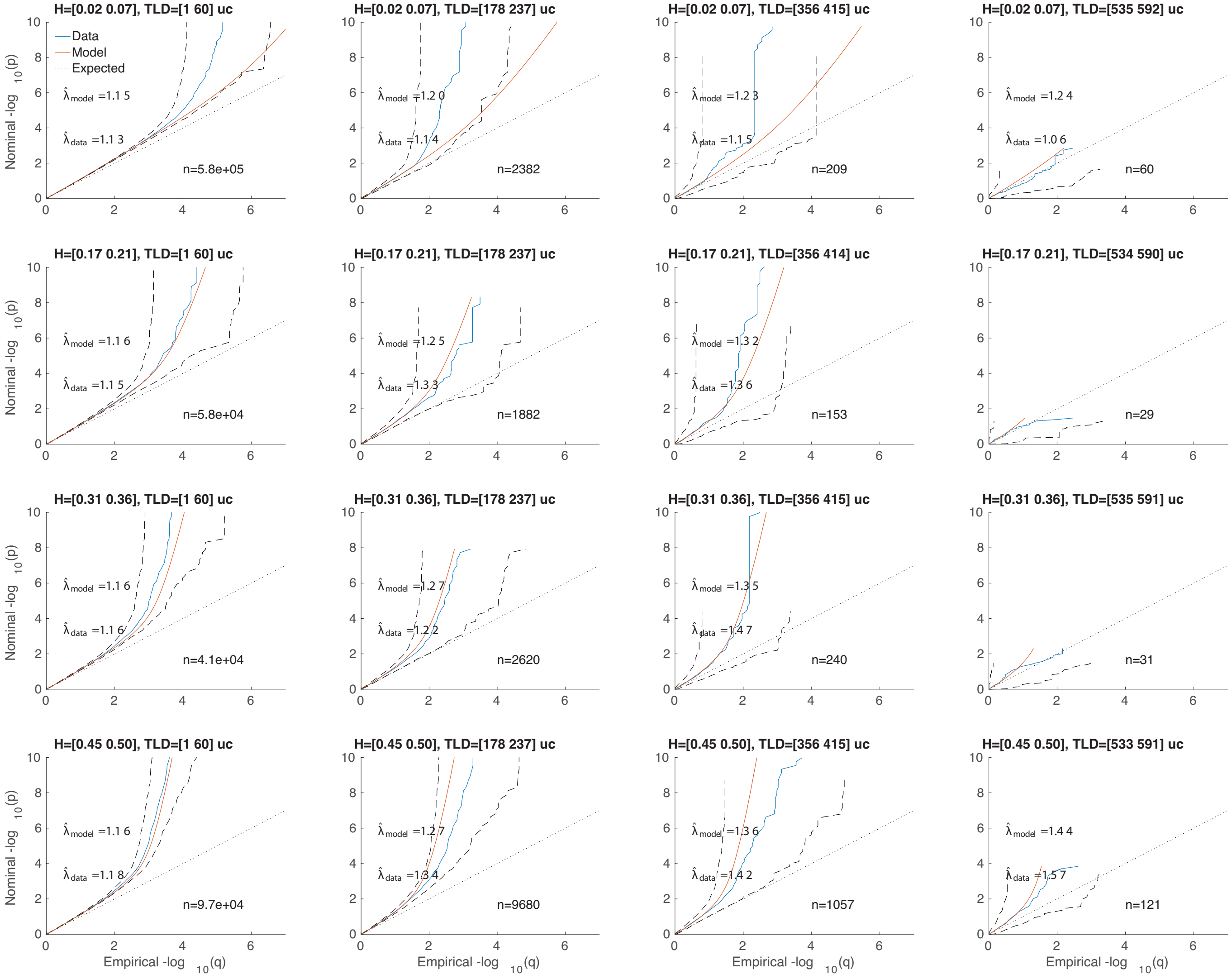


Figure S15: A 4×4 heterozygosity×total-LD grid of QQ plots for ulcerative colitis. See Figures 1 (D) and S12.

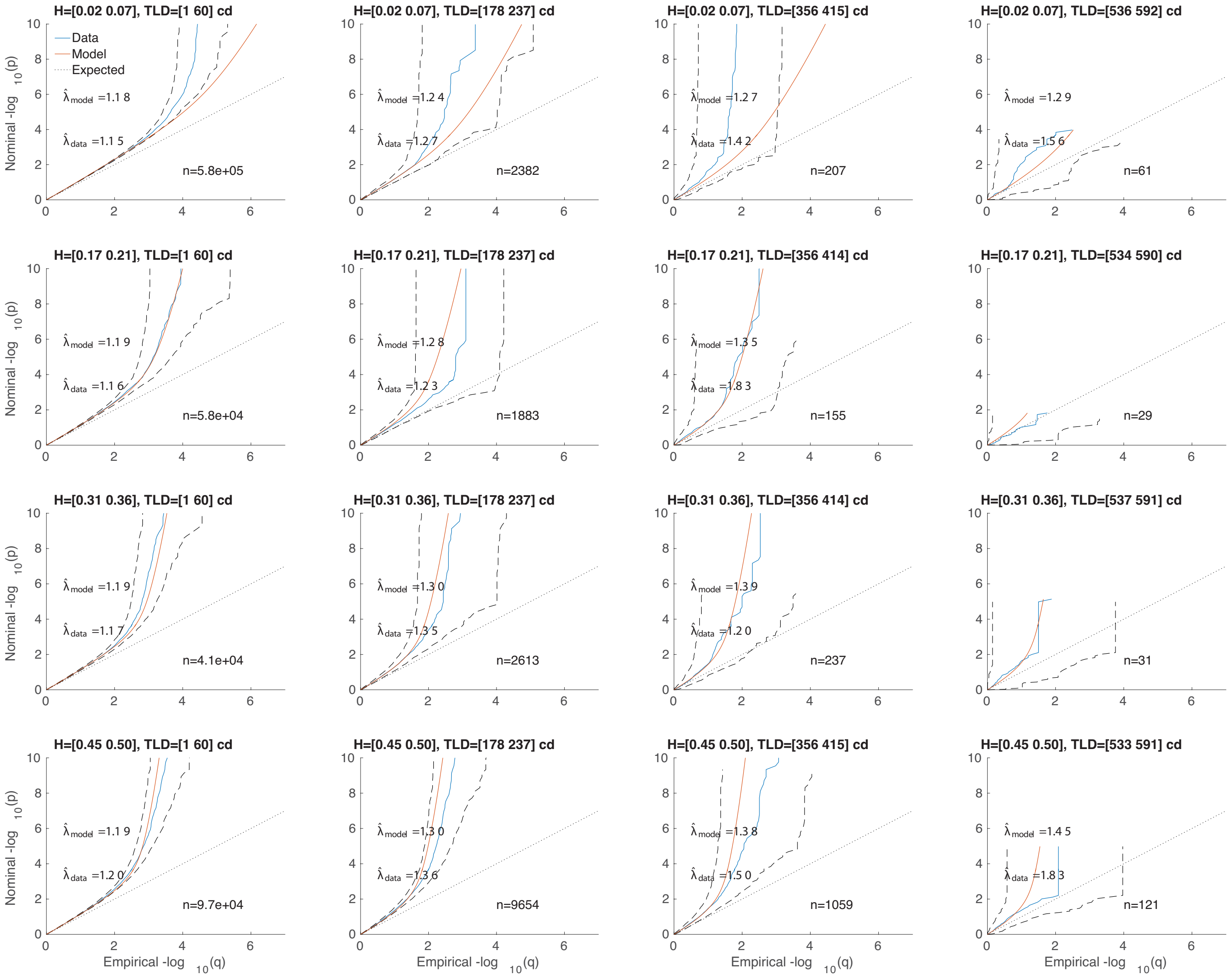

Figure S16: A 4×4 heterozygosity×total-LD grid of QQ plots for Crohn's disease. See Figures 1 (E) and S12.

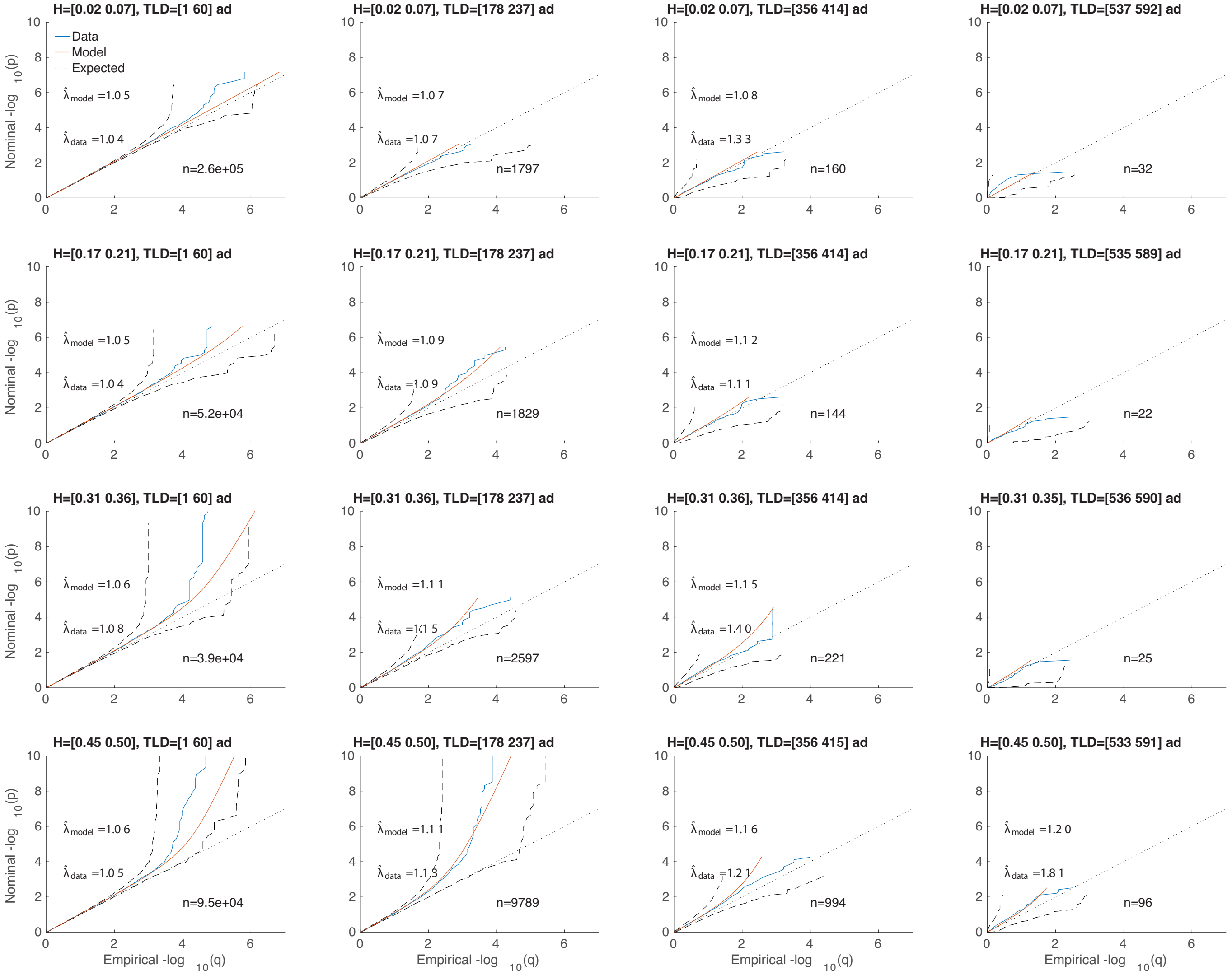


Figure S17: A 4×4 heterozygosity×total-LD grid of QQ plots for Alzheimer's disease, excluding chromosome 19. See Figures 1 (F), S7, S12, and S18.

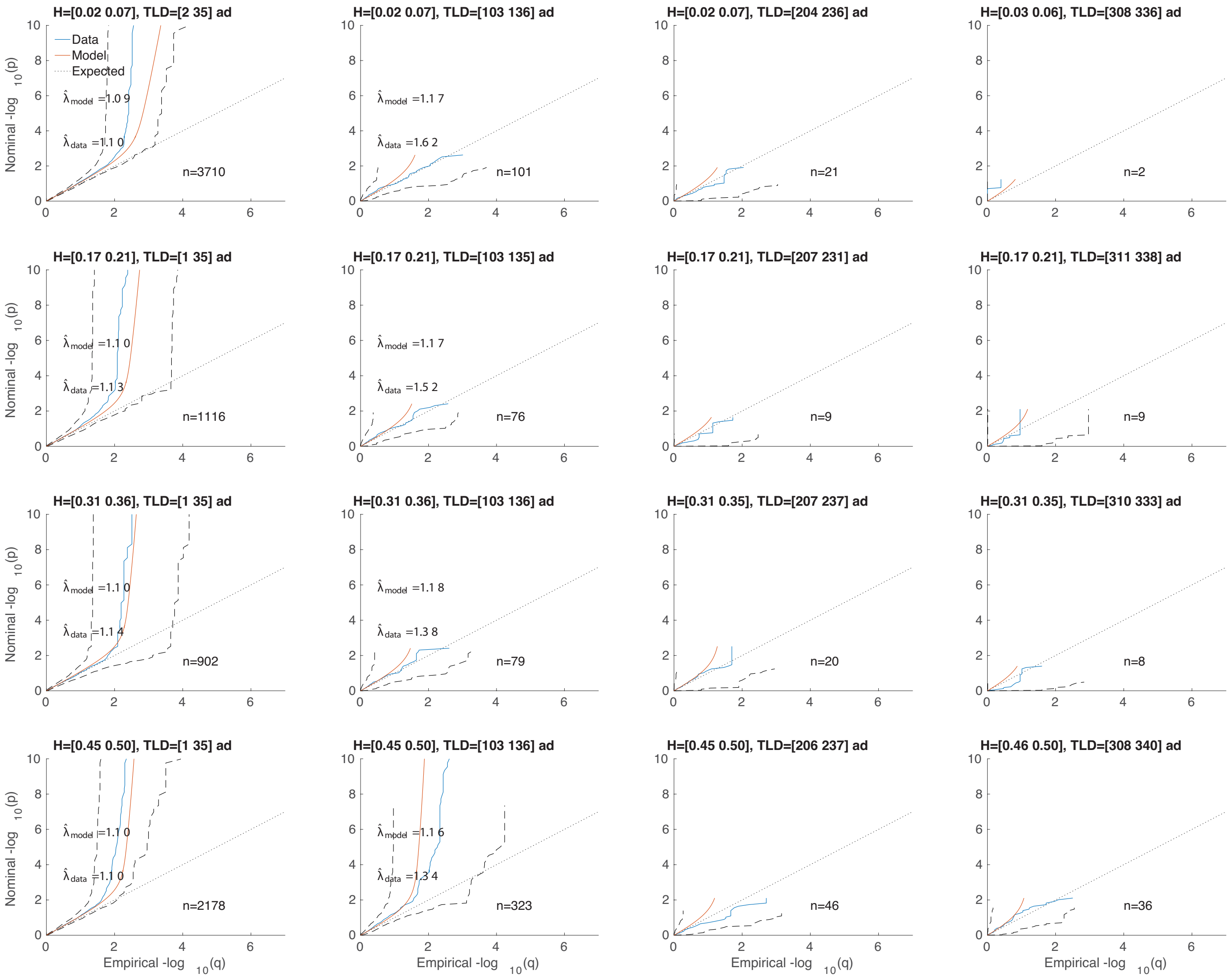


Figure S18: A 4×4 heterozygosity×total-LD grid of QQ plots for Alzheimer's disease, chromosome 19 only. See Figures 1 (F), S7, S12, and S17.

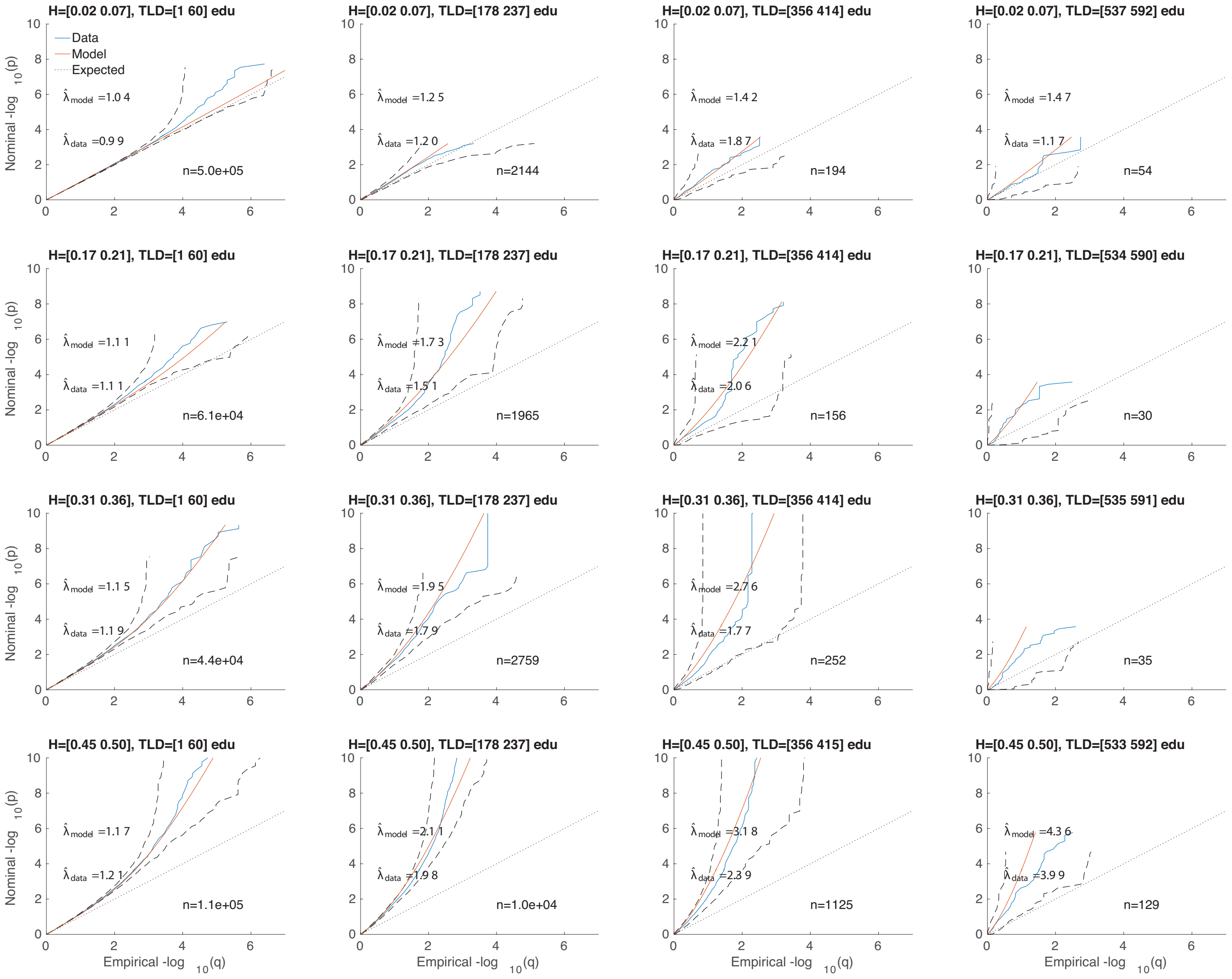

Figure S19: A 4×4 heterozygosity×total-LD grid of QQ plots for years of education. See Figures 2 (A) and S12.

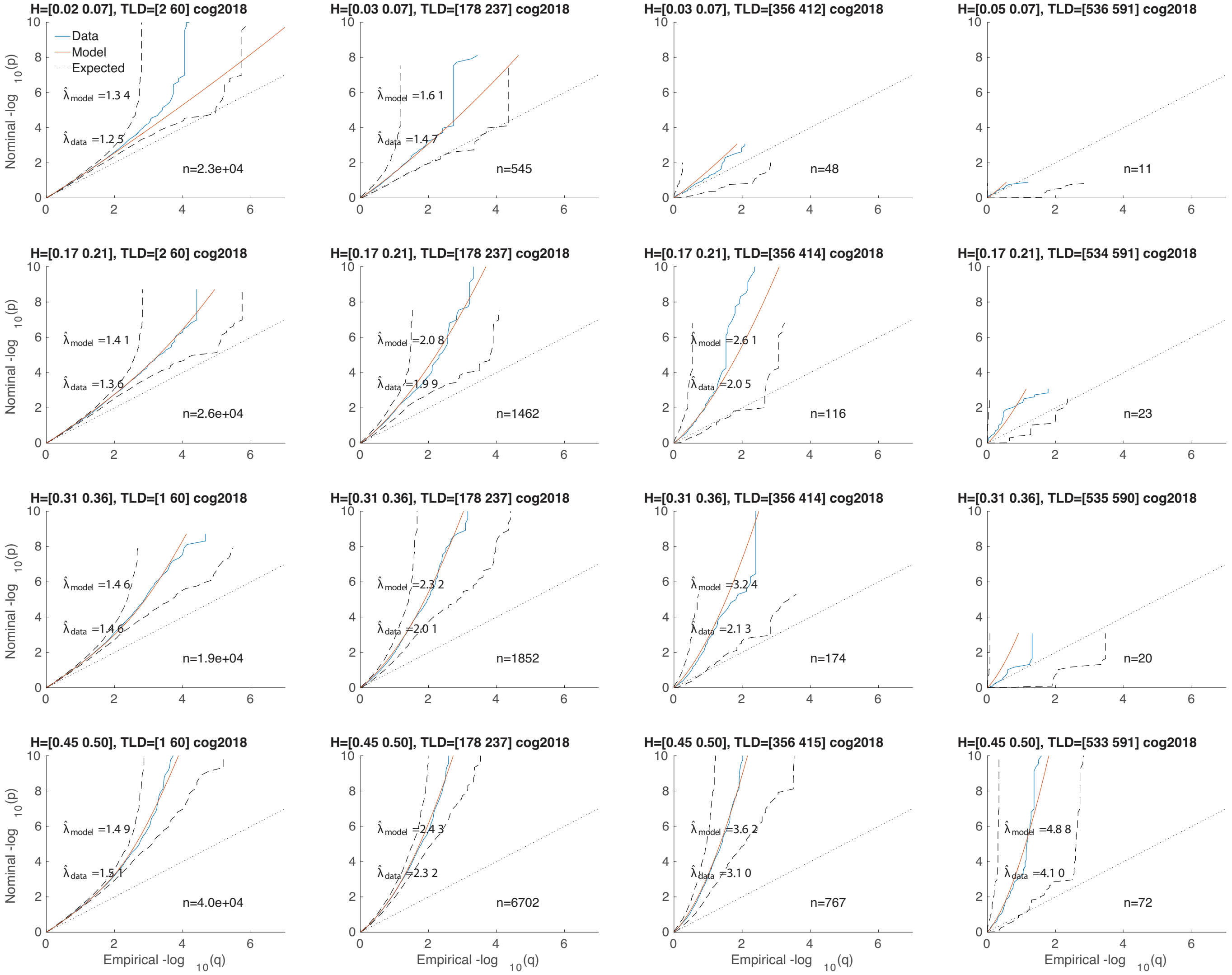


Figure S20: A 4×4 heterozygosity×total-LD grid of QQ plots for intelligence. See Figures 2 (B) and S12.

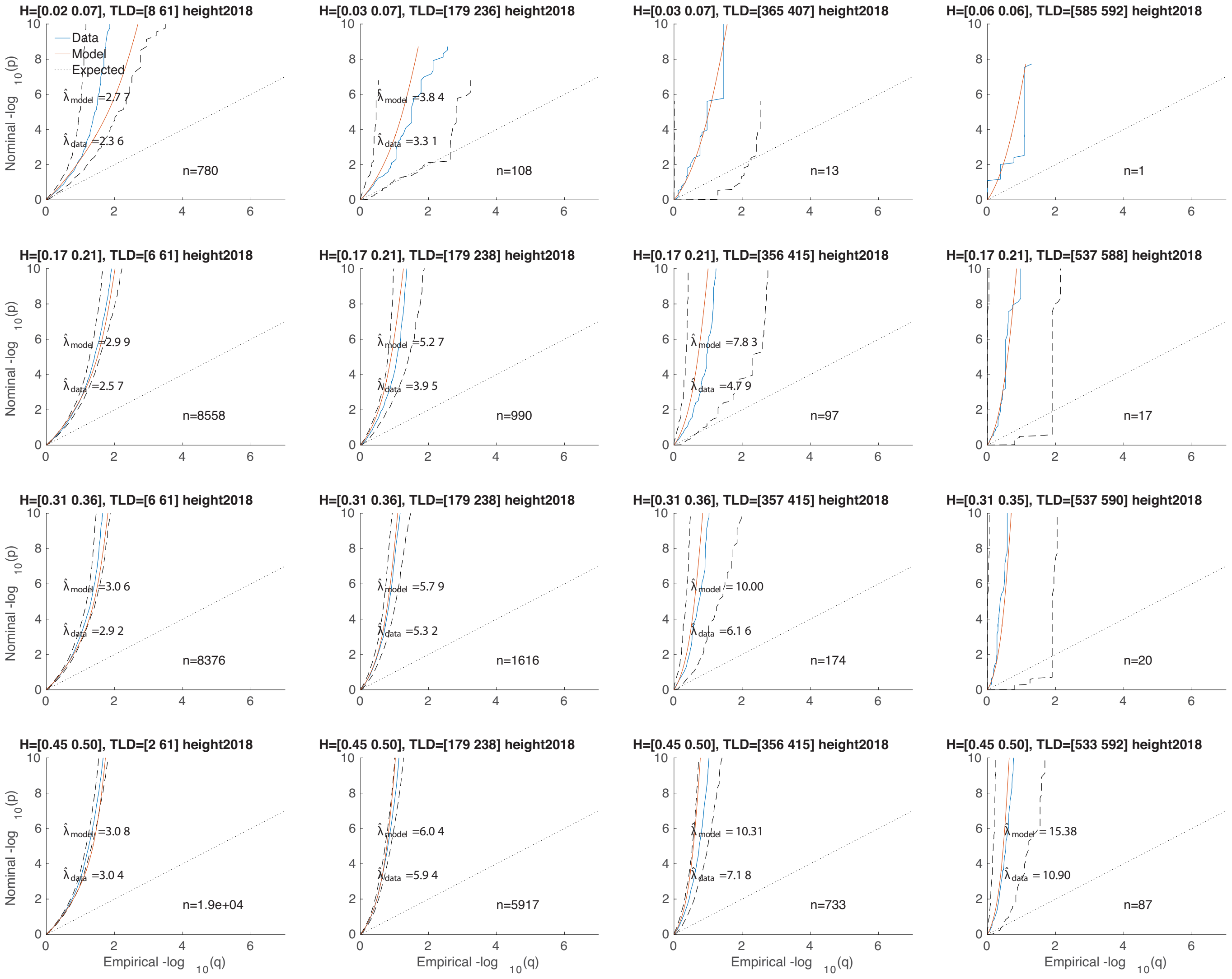

Figure S21: A 4×4 heterozygosity×total-LD grid of QQ plots for height. See Figures 2 (C) and S12.

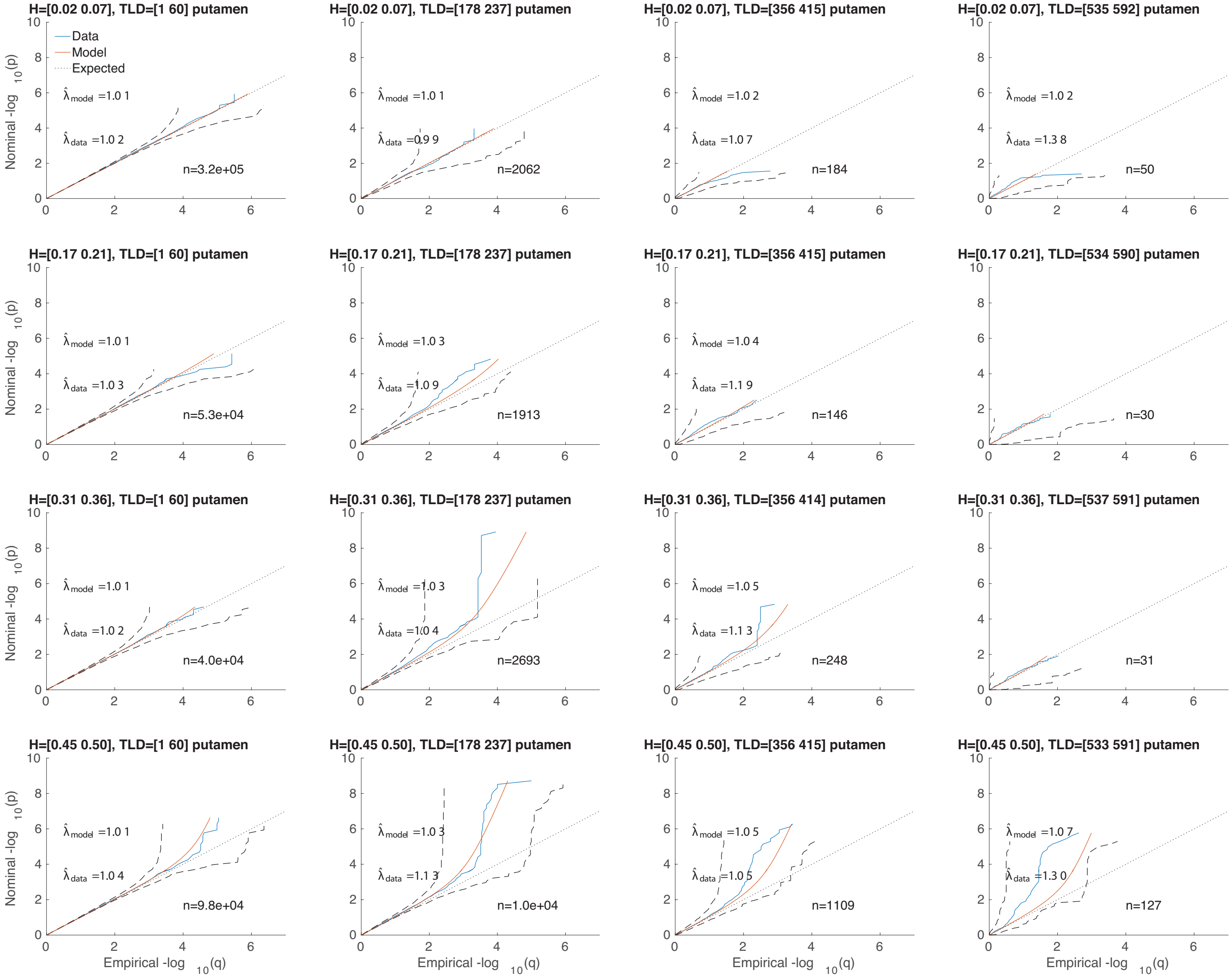


Figure S22: A 4×4 heterozygosity×total-LD grid of QQ plots for putamen volume. See Figures 2 (D) and S12.

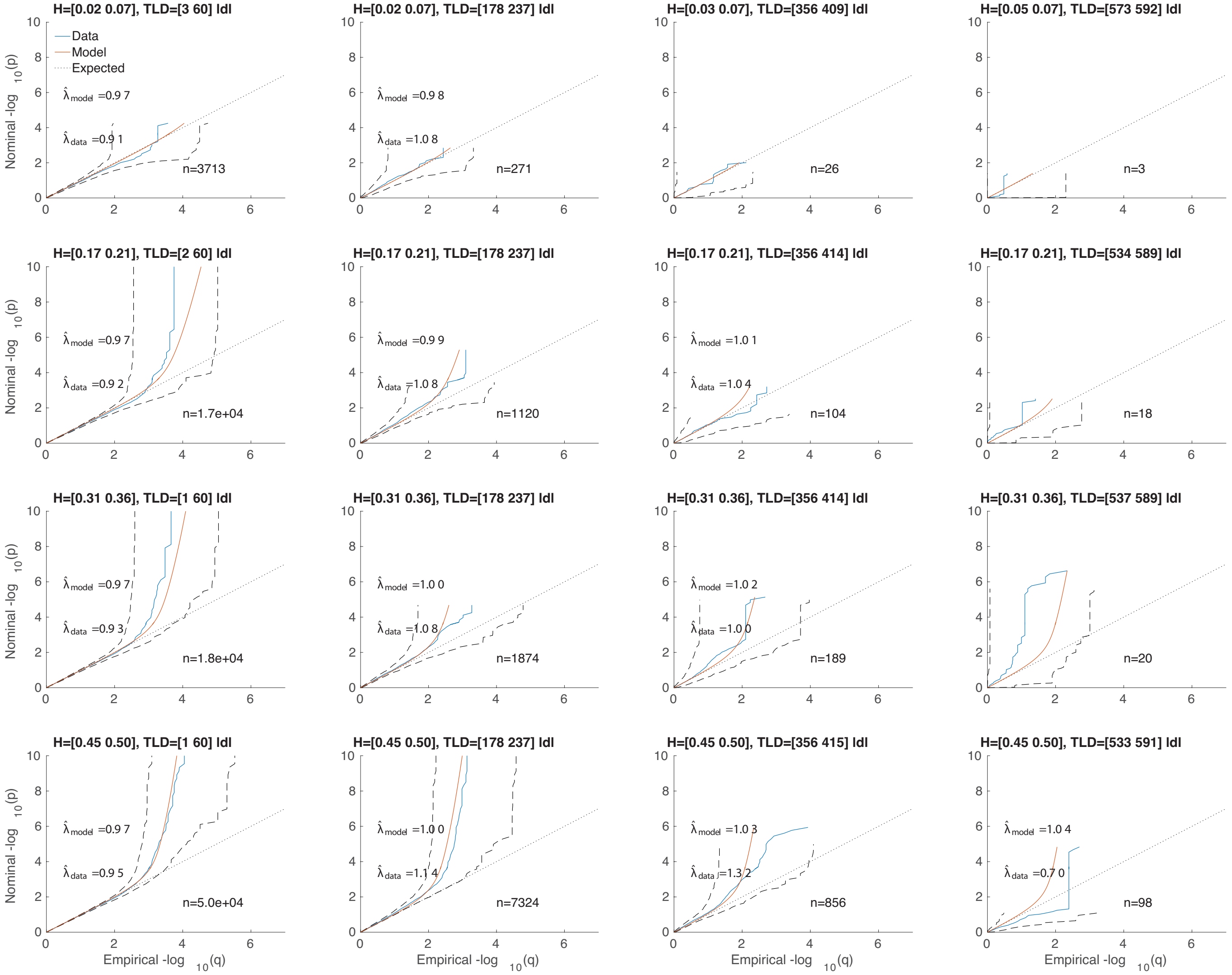


Figure S23: A 4×4 heterozygosity×total-LD grid of QQ plots for low density lipoprotein. See Figures 2 (E) and S12.

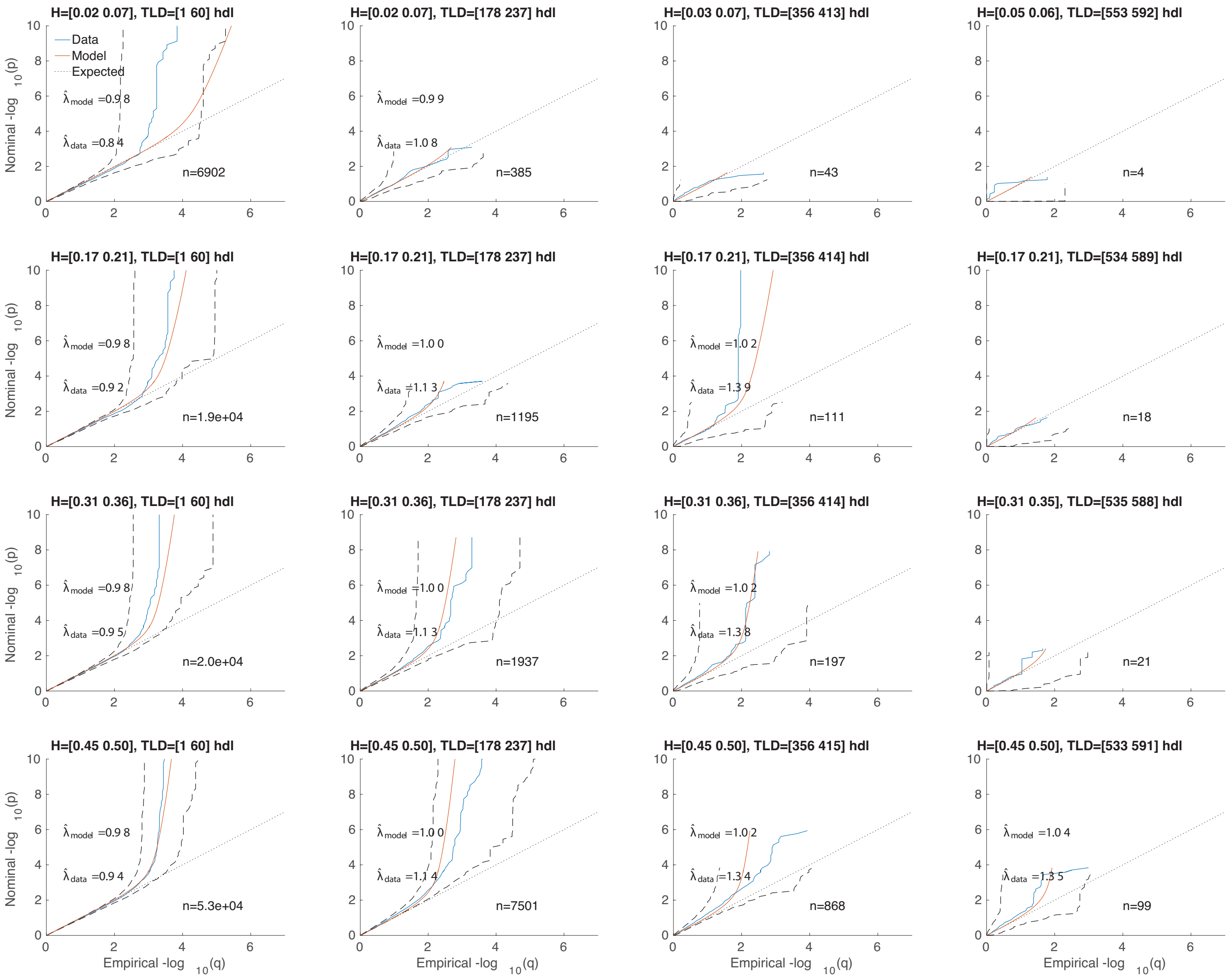

Figure S24: A 4×4 heterozygosity×total-LD grid of QQ plots for high density lipoprotein. See Figures 2 (F) and S12.

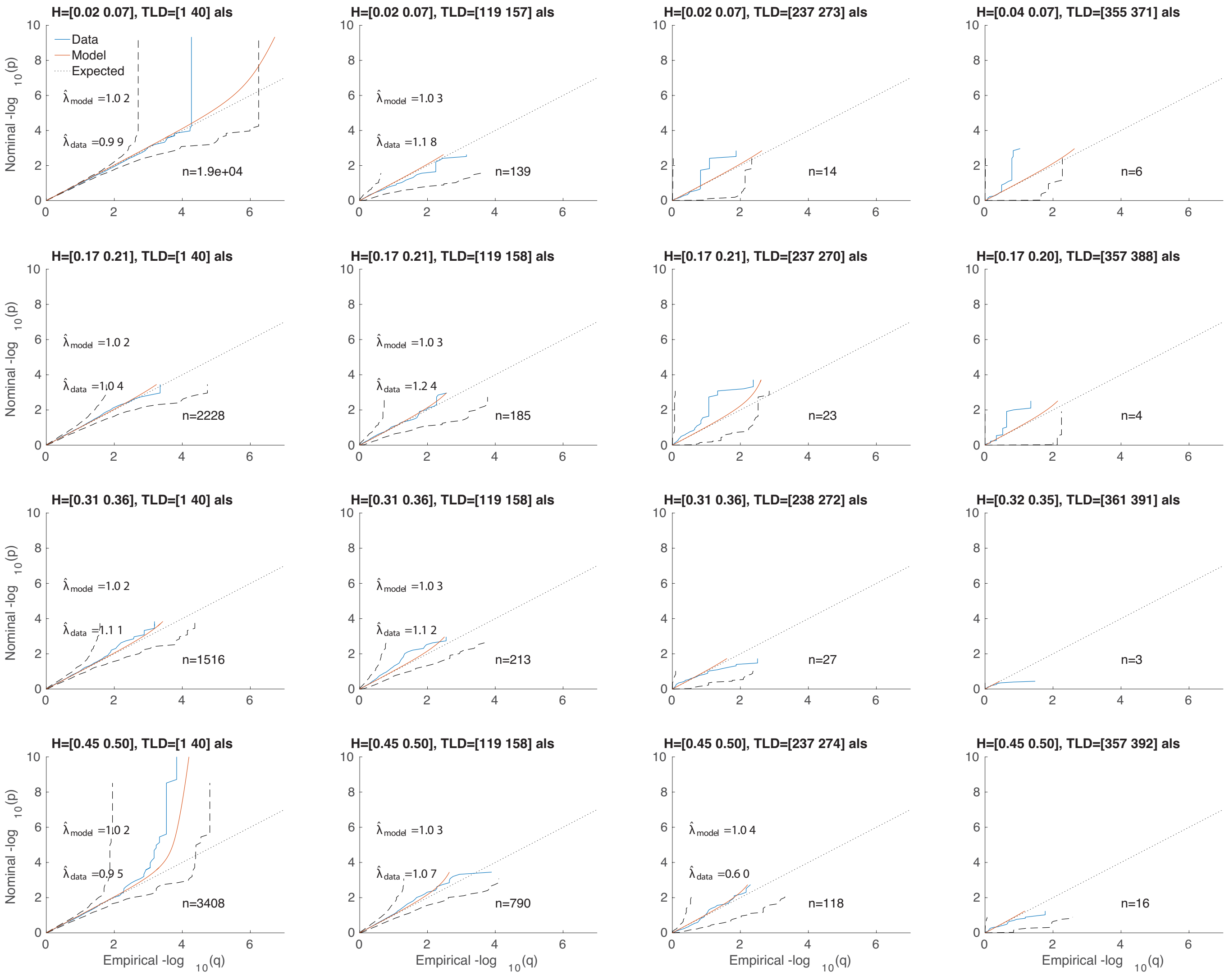


Figure S25: A 4×4 heterozygosity×total-LD grid of QQ plots for amyotrophic lateral sclerosis, chromosome 9 only. See Figures 1 (H) and S12.

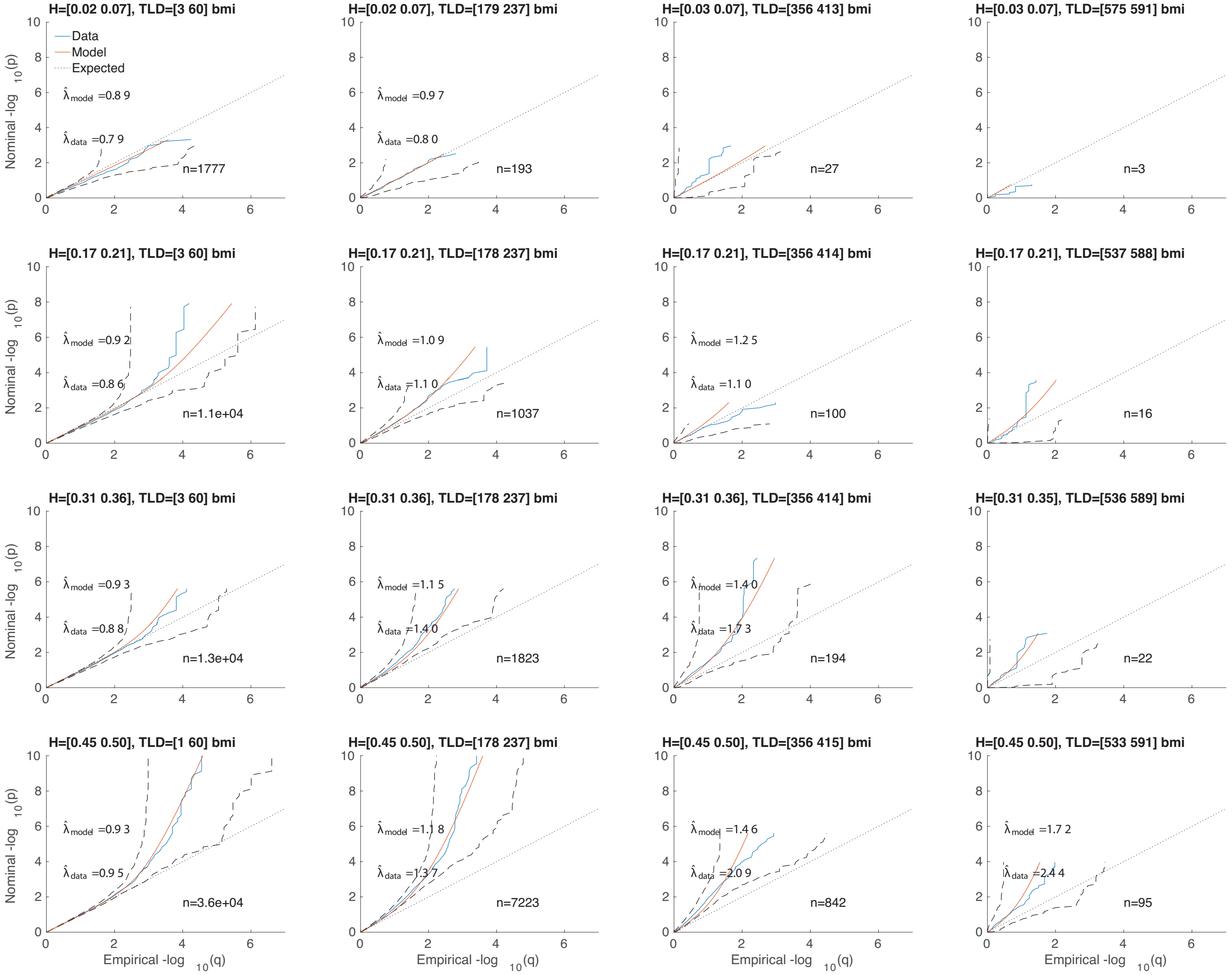


Figure S26: A 4×4 heterozygosity×total-LD grid of QQ plots for body mass index. See Figures 1 (C) and S12.

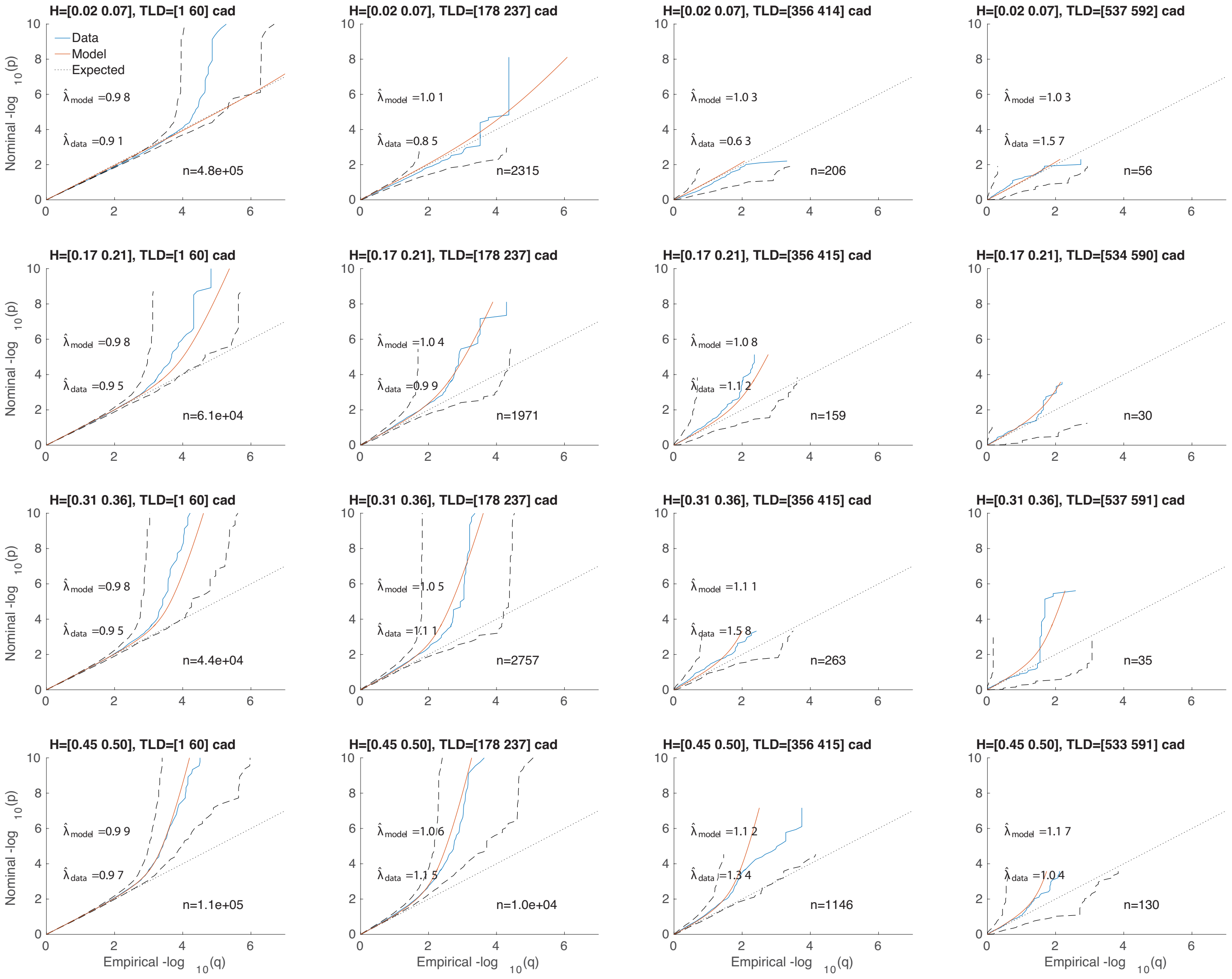

Figure S27: A 4×4 heterozygosity×total-LD grid of QQ plots for coronary artery disease. See Figures 1 (D) and S12.

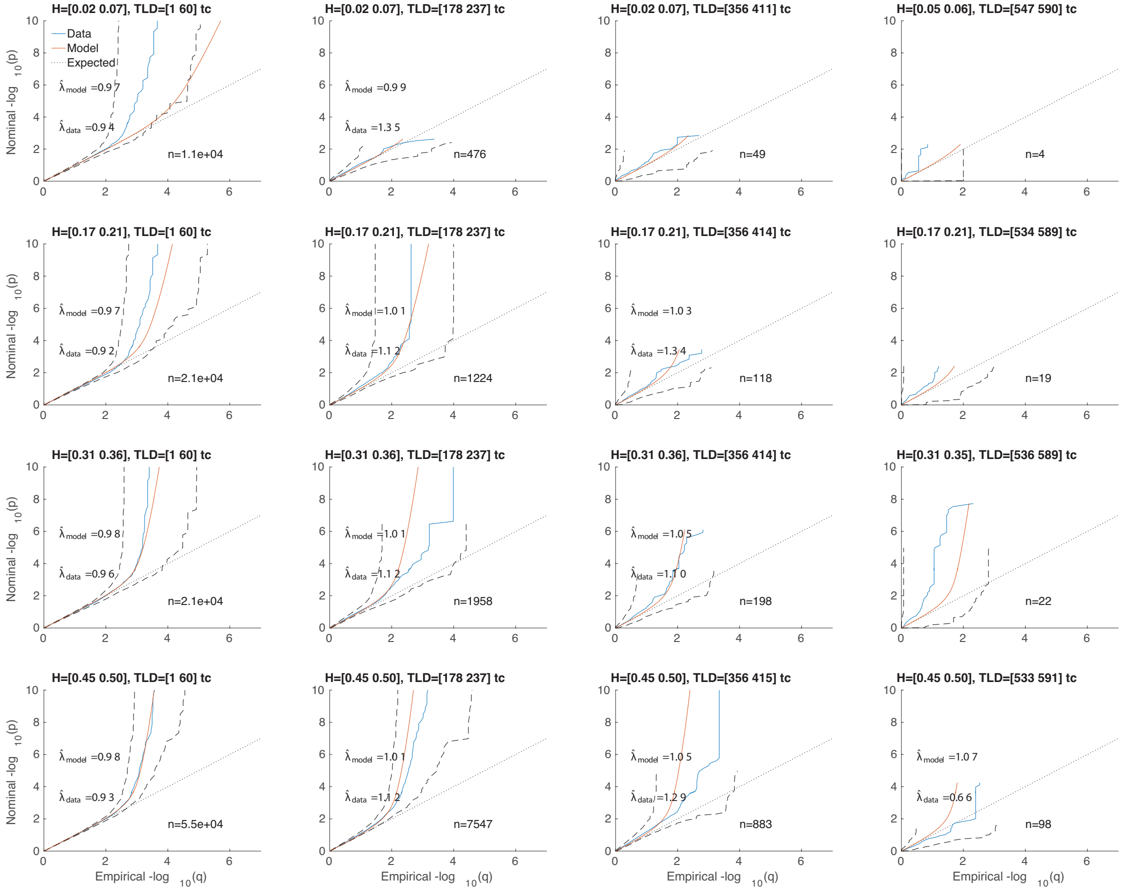

Figure S28: A 4×4 heterozygosity×total-LD grid of QQ plots for total cholesterol. See Figures 1 (H) and S12.
